# PTPRJ-Targeting Peptide Agonist Induces Broad Cellular Signaling Perturbations and DNA Damage in Lung Cancer Cells

**DOI:** 10.1101/2025.11.17.688709

**Authors:** Sophie Rizzo, William S. Hart, Rachel L. Fetter, Bokai Song, Forest M. White, Matthew J. Lazzara, Damien Thévenin

**Affiliations:** Department of Chemistry, Lehigh University, Bethlehem, PA, USA; Department of Chemical Engineering, University of Virginia, Charlottesville, VA, USA; Department of Biological Engineering, Massachusetts Institute of Technology, Cambridge, MA, USA; Koch Institute for Integrative Cancer Research, Massachusetts Institute of Technology, Cambridge, MA, USA

## Abstract

Receptor protein tyrosine phosphatases (RPTPs) are key regulators of cell signaling. However, their study and therapeutic targeting have been limited by the lack of known natural ligands or selective agonists, as well as an incomplete understanding of their structure-function relationships. Nonetheless, receptor homodimerization has been shown to suppress RPTP catalytic activity by restricting substrate access, offering a promising strategy for examining and modulating their function. Our previous work on PTPRJ, a member of the RPTP family, showed that its transmembrane domain regulates homodimerization, thereby controlling access to receptor tyrosine kinase (RTK) substrates and their phosphorylation levels. We also developed peptides that disrupt this dimerization, thereby inhibiting RTK phosphorylation and reducing cancer cell migration. These peptides were then engineered for selective, pH-sensitive insertion into the acidic tumor microenvironment to enhance efficacy while limiting off-target effects. Yet how broadly PTPRJ activation reshapes the phosphotyrosine landscape and whether those changes yield coherent cellular phenotypes remains unclear. In this study, we employed tyrosine phosphoproteomics, immunoblotting, immunofluorescence, and functional assays to assess the global impact of our lead peptide candidate, Hybrid 7, in A549 lung cancer cells that endogenously express PTPRJ. We find that Hybrid 7 decreases EGFR phosphorylation and selectively reduces phosphorylation across additional RTKs and motility adaptors, producing strong inhibition of EGF-driven migration and reduced proliferation. Hybrid 7 also elevates reactive oxygen species and DNA damage, and enforces CDK1-dependent G2/M arrest, indicating a primarily cytostatic, checkpoint-mediated response. These findings highlight the potential of RPTP-targeting peptides as valuable tools for dissecting RPTP function and as possible therapeutic agents capable of modulating key oncogenic pathways and inhibiting cancer progression.

## Introduction

Protein tyrosine phosphorylation is crucial for the initiation and propagation of cell signaling transduction in response to external stimuli.^1,2^ The timing and duration of phosphorylation are tightly controlled through the counterbalancing actions of receptor tyrosine kinases (RTKs) and receptor protein tyrosine phosphatases (RPTPs).^3^ RPTPs modulate the signal-initiating potency of RTKs by dephosphorylating key regulatory tyrosine residues.^4–6^ It is well understood that the disruption of RTK and RPTP activity through oncogenic mutations can lead to unregulated protein tyrosine phosphorylation, which is linked to many forms of cancer.^7–11^ Therapies have been designed to target oncogenic RTK signaling, such as that initiated by the epidermal growth factor receptor (EGFR). Cetuximab and gefitinib are two examples of highly effective treatments that target EGFR. However, gatekeeper mutations and the bypassing of signaling through other RTKs make these treatments prone to acquired resistance mechanisms that can affect their long-term efficacy.^12–16^

RPTPs have recently been recognized as promising therapeutic targets because they can attenuate RTK signaling and suppress the growth of multiple tumor types, including lung, colon, thyroid, and breast cancers.^17–21^ Therefore, enhancing phosphatase activity toward RTKs could offer an alternative strategy to inhibit oncogenic RTK signaling and malignant phenotypes while avoiding the common resistance mechanisms. However, the development of agonists and antagonists for RPTPs is still limited due to an insufficient understanding of their structure-function relationship.^22–27^

Like RTKs, RPTP homodimerization has been shown to be stabilized through transmembrane (TM) domain interactions.^28–31^ However, unlike RTKs, homodimerization appears to inhibit the activity of RPTPs, potentially due to restricted access to the catalytic domain in the dimer form.^28,32,33^ We previously reported that specific TM domain interactions stabilize the homodimerization of a member of the RPTP family, protein tyrosine receptor J (PTPRJ). PTPRJ, like many RPTPs, is a tumor suppressor and regulator of growth factor signaling, including that by PDGFβR, VEGFR, METR, FLT3, and EGFR.^34–43^ We previously identified three glycine residues (G979, G983, and G989) as major mediators of PTPRJ homodimerization through a specific contact interface. Introducing glycine-to-leucine (small-to-large) mutations disrupted PTPRJ oligomerizations in cells, promoted PTPRJ association with EGFR, decreased EGFR phosphorylation (Y1068 and Y1173), and inhibited EGFR-driven phenotypes.^29^ Similar results were observed with the FLT3 kinase receptor (containing oncogenic internal tandem duplications) in acute myeloid leukemia cell models.^44^ Thus, disrupting TM-mediated oligomerization could provide a strategy to modulate RPTP activity and, consequently, RTK signaling in cells.

To that end, we previously designed a TM peptide, RJ_binder_, that can interact with the TM domain of PTPRJ and disrupt its TM domain-mediated homodimerization. In EGFR-driven cancer cells, RJ_binder_ decreased PTPRJ self-association, enhanced PTPRJ interactions with EGFR, inhibited EGFR phosphorylation, and suppressed cell migration.^29^ Although this peptide is the first RPTP agonist of its kind, it has limited solubility in water due to its transmembrane nature. Additionally, it requires a delivery vehicle, such as detergent micelles, and fails to insert unidirectionally into membranes. Moreover, it embeds indiscriminately in both healthy and cancerous cells, which restricts its effectiveness and potential uses. To address these limitations and better target RTK-dysregulated cancers, we combined the properties of RJ_binder_ with the pH (Low) Insertion Peptide (pHLIP), another TM peptide that inserts into cancer cell membranes by exploiting the inherent acidic microenvironment of solid tumors.^45–50^ Doing so produced hybrid peptides; among them, Hybrid 7 is water-soluble, inserts as a helical peptide at mildly acidic pH mimicking the microenvironment of tumors, and functions as a PTPRJ TM agonist.^51^ In human squamous cell carcinoma HSC3 cells engineered to overexpress PTPRJ, Hybrid 7 increased PTPRJ activity toward EGFR, resulting in a significant reduction in EGFR phosphorylation, migration, and proliferation. In contrast, no effect was observed in parental HSC3 cells lacking PTPRJ. These results indicate that Hybrid 7 can modulate EGFR and related phenotypes.

However, the systems-level consequences of PTPRJ activation by Hybrid 7 on phosphotyrosine signaling and cellular phenotypes remain undefined. To address this, we integrated tyrosine phosphoproteomics with immunoblotting, immunofluorescence, and functional assays in this study to quantify the global effects of Hybrid 7 in A549 lung cancer cells that endogenously express PTPRJ. We hypothesized that the response to peptide treatment would be context-dependent, influenced by factors such as cell type, RTK expression profiles, the autoregulatory behavior of PTPRJ, and the baseline activity of intracellular signaling networks. We found that Hybrid 7 reduces the phosphorylation of EGFR, additional RTKs, and motility adaptors, blocking EGF-driven migration and slowing proliferation. It also elevates reactive oxygen species and DNA damage, and enforces CDK1-dependent G2/M arrest, indicating a cytostatic, checkpoint-mediated response and supporting RPTP-targeting peptides as potential therapeutic tools.

## Results

### Hybrid 7 inhibits EGFR and EGFR-driven phenotypes in A549 Cells

Based on transcriptomic and proteomic datasets from the Cancer Cell Line Encyclopedia (DepMap Public 25Q3; https://depmap.org/portal), we selected A549 as a cell model for this study because of its relatively high endogenous PTPRJ and wild-type EGFR expression levels (**Fig. S1**).^52^ Treating A549 cells with Hybrid 7 (10 µM for 10 minutes) reduced EGFR phosphorylation (by about 40%) in a pH-dependent manner (**Fig. 1a, b**). Hybrid 7 also inhibits cell viability in a pH-dependent manner, with an IC_50_ value of approximately 1.4 µM (**Fig. 1c**).^51^ It is important to note that cells are exposed to low-pH media for only 10 minutes before being recovered in normal culture media, and that the decrease in cell viability is not due to the low pH treatment. Immunostaining of Ki-67, a protein expressed in cells that are actively dividing, confirms that Hybrid 7 inhibited cell proliferation (**Fig. 1d, e**). Hybrid 7 also inhibited wound closure in a pH-dependent manner, as we observed previously in HSC3^+^ cells (**Fig. 1f, g**). Interestingly, while the inhibition of cell migration is only observed in the presence of EGF (**Fig. 1f, g**), cell proliferation is inhibited under both unstimulated and EGF-stimulated conditions, suggesting that EGFR activation may not be necessary for Hybrid 7 to exert its antiproliferative effects.

**Figure 1.**
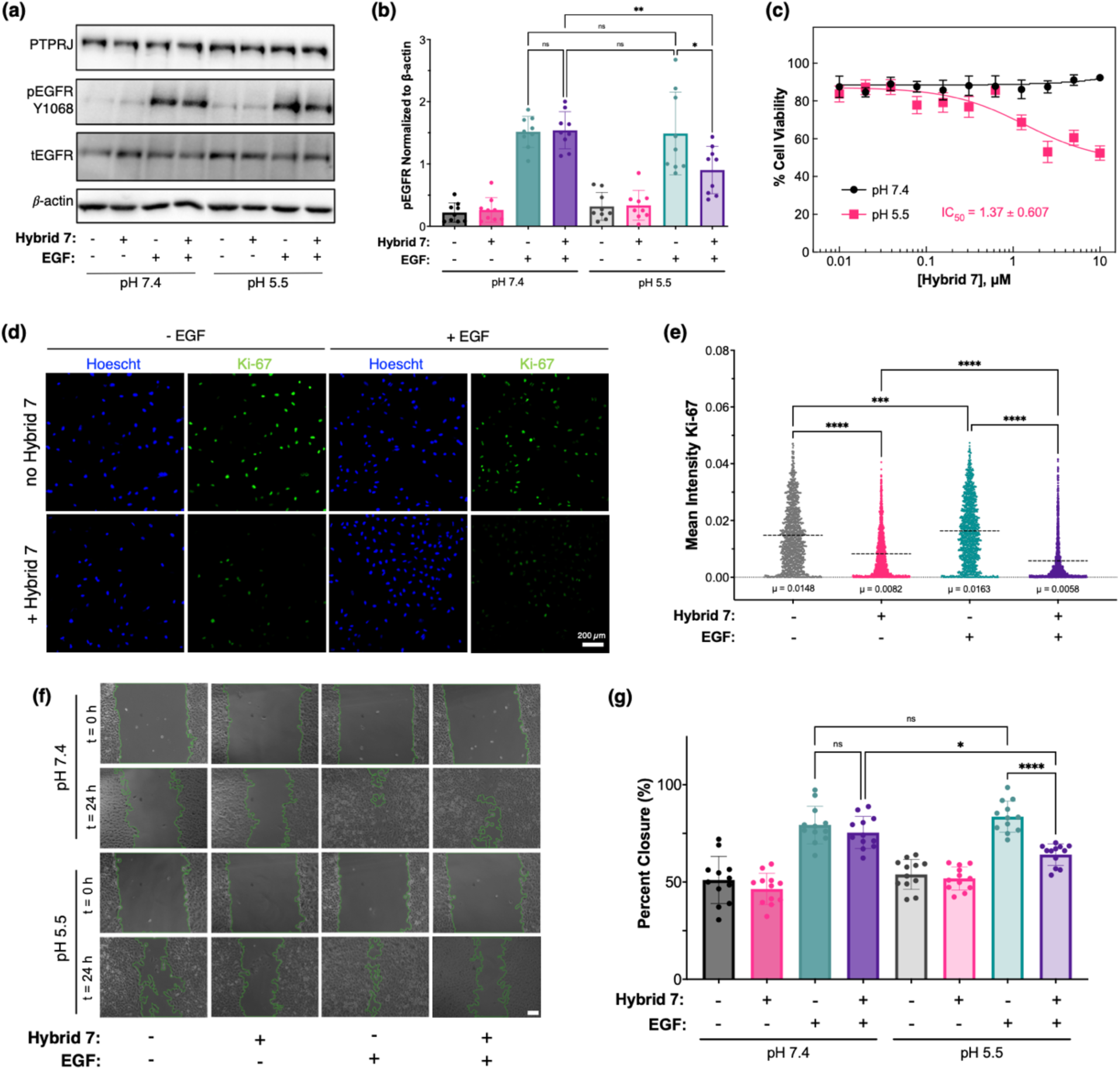
Hybrid 7 inhibits EGFR phosphorylation and A549 cell proliferation. **(a)** Representative immunoblot of total and phospho-protein levels after treatment with +/- Hybrid 7 (10 µM for 10 min) and +/- EGF (10 ng/mL for 10 min). (**b)** Quantification of pEGFR (Y1068). Results are shown as mean ± S.D. (n = 9). **(c)** Effect of Hybrid 7 (10 µM for 10 min) on cell viability measured by the colorimetric MTT assay after 72 hours. All measurements are normalized to cells treated at pH 7.4 with no Hybrid 7 (100% viability). Results are shown as mean ± S.D. (n = 3). Data were fitted to a non-linear model to determine the IC_50_ value. **(d)** Representative immunofluorescence images of the effect of Hybrid 7 (10 µM for 10 min at pH 5.5) treatment on Ki-67 proliferation marker determined by immunofluorescence after 18 hours. Scale bar **(e)** Quantification of mean Ki-67 stain intensity within the nucleus. **(f)** Representative phase-contrast images with tracing to identify open scratch areas (Scale bar: 200 µM) and **(g)** quantification of the effect of Hybrid 7 on wound closure. A549 cells were starved for 18 hours in 1% FBS media before treatment with 10 µM Hybrid 7 at the indicated pH, scratched (t = 0h), and incubated without EGF or with EGF (10 ng/mL) until the wound closed. Relative closure was quantified by calculating the percent change in area between the 0 h and 24 h images. Results are shown as mean ± S.D. (n = 12). The statistical significance of differences between samples was determined by one-way ANOVA with Tukey’s multiple comparisons correction (α = 0.05). ****, p ≤ 0.0001; ***, p ≤ 0.001; *, p ≤ 0.05; ns, p > 0.05. Only a subset of comparisons is shown for clarity.

Finally, although Hybrid 7 significantly affects EGFR phosphorylation and EGFR-driven cell migration and proliferation in A549 cells, Luminex multiplex analysis showed no notable changes in key downstream signaling molecules, such as Akt, ERK1/2, and others involved in the Akt, mTOR, MAPK, and SAPK pathways, under both unstimulated and EGF-stimulated conditions upon Hybrid 7 treatment (**Fig. S2**). These data suggest that while Hybrid 7 affects EGFR phosphorylation and activity (via the disruption of PTPRJ dimer), its mechanism of action may result in many relatively small changes that are difficult to detect or extend beyond the pathways measured by Luminex. Indeed, as a PTPRJ agonist, Hybrid 7 could modulate a wide array of PTPRJ substrates beyond EGFR that may contribute to the observed phenotypic effects.

### Global analysis of Hybrid 7 on phospho-tyrosine cell signaling by mass spectrometry

To better understand the overall effects of Hybrid 7 treatment on signaling networks and to potentially identify the pathways responsible for the observed phenotypic results, we used mass spectrometry-based phosphotyrosine (pTyr) proteomics. Accordingly, we focused our efforts on low-pH treatments because they lead to the active, inserted conformation of Hybrid 7. Results show that a total of 209 phosphorylated proteins with 343 unique pTyr sites were identified and quantified from this dataset (**Fig. 2 and Table S1**).

**Figure 2.**
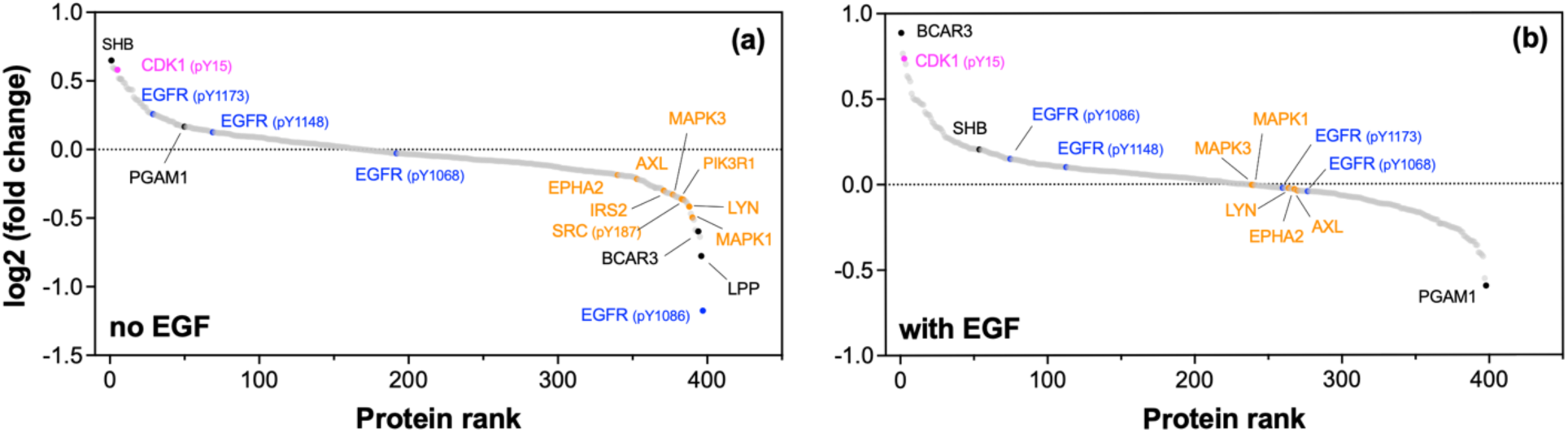
Ranked abundance plot of all phosphorylated proteins detected in the study. Proteins are ranked by their log2 fold change (+Hybrid 7 / no Hybrid 7) in the absence or presence of EGF (10 ng/mL for 10 min). Positive log2 fold change values represent pTyr hyperphosphorylation upon Hybrid 7 treatment (n=3).

These results raise several key observations. First, Hybrid 7 treatment leads to changes in phosphorylation at key signaling molecules, especially under unstimulated conditions (**Fig. 2**). Most notably, Hybrid 7 treatment led to over 50% decrease in EGFR phosphorylation at position Y1086 (**Fig. 2a**). Importantly, phosphorylated Y1086 binds the growth factor receptor-bound protein 2 (Grb2) adaptor protein, which activate Erk MAPK pathway and thus mediate cell migration and proliferation phenotypes.^53–58^ Several EGFR-related signaling molecules were also identified as hypo-phosphorylated upon peptide treatment, including MAPK1 (Y187), Lyn (Y194), PIK3R1/p85a (Y467), Src (Y187), and a few RTKs such as EphA2 (Y575, Y587, Y772) and Axl (Y452) (**Fig. 2a**). Interestingly, other proteins whose Tyr phosphorylation promotes cancer cell migration, invasion, or adhesion are also found to be hypo-phosphorylated upon treatment with Hybrid 7, including LPP (Lipoma Preferred Partner),^59^ BCAR3 (Breast Cancer Anti-Estrogen Resistance 3),^60–67^ and PGAM1 (Phosphoglycerate Mutase 1).^68–71^ While many of these downstream proteins do not have a clear link to EGFR receptor phosphorylation, they are linked to downstream signaling pathways that coincide with EGFR and other tyrosine-mediated signaling pathways, such as Src-family kinases, Cas, and Ras, that play critical roles in propagating cancer phenotypes. On the other hand, EGFR phosphorylation at other detected EGFR pY sites (1068, 1148, and 1173) shows either a slight increase or decrease in phosphorylation under unstimulated or stimulated conditions. The 1148 pY site is among the most abundant phosphorylation sites upon EGFR autophosphorylation, and similar to the Y1173 site, is involved in providing a docking site for other adaptor proteins, including Ras/MAPK, Shc, SHP-1, and PLC𝛾 , which contribute to the effect on EGFR-mediated mitogenesis and transformation.^54,72–79^ Notably, phosphorylation at EGFR pY1173 is reduced under stimulated conditions when compared to unstimulated conditions, while phosphorylation at pY1148 stays relatively the same in both conditions. Still, phosphorylation at Y1068 is somewhat reduced under stimulated conditions (**Fig. 2b**), which is consistent (albeit not in amplitude) with the immunoblotting results (**Fig. 1a, b**).

Second, Hybrid 7 exhibits weaker effects under EGF stimulation compared to unstimulated cells. These diminished responses, evident for MAPK1, Lyn, PIK3R1, and EphA2 (**Fig. 2b**), are likely due to EGF-induced phosphorylation masking the subtler changes caused by Hybrid 7. It is also possible that this single time point (10 min post-EGF stimulation) may limit the detection of specific phosphorylation events occurring at other time points. Third, while other proteins are shown to be more hyper- or hypo-phosphorylated depending on the conditions (e.g., SHB, BCAR3, and PGAM1), Hybrid 7 consistently enhances CDK1 phosphorylation at Tyr15 (by more than 50% under unstimulated and by more than 70% under stimulated conditions), making CDK1 unique in the dataset. CDK1 is a critical regulator of the G2/M phase transition, an essential checkpoint in the cell cycle that ensures replication is complete and that DNA is undamaged before mitosis begins. CDK1 is a critical regulator of the G2/M transition, and its activity is tightly controlled by phosphorylation. Specifically, WEE1-mediated phosphorylation of Y15 inhibits CDK1 and cell-cycle progression. When sustained, CDK1 hyperphosphorylation leads to G2 arrest, accumulation of DNA damage, and ultimately cell death. Therefore, we hypothesize that Hybrid 7 attenuates EGFR/RTK signaling, affecting not only cell proliferation and migration but also compromising the cell cycle and DNA damage response.

### CDK1 hyperphosphorylation is linked to growth suppression

First, to confirm that elevated CDK1 phosphorylation results from Hybrid 7 treatment and establish its central in the anti-proliferative effect of Hybrid 7, we hypothesized that inhibiting CDK1 phosphorylation (using the potent WEE-1 inhibitor MK-1775) before treating with Hybrid 7 would not only reduce the CDK1 hyperphosphorylation caused by Hybrid 7 but also restore cell proliferation. Immunoblotting shows that Hybrid 7 treatment resulted in an increase in phosphorylation of CDK1 Y15 under both unstimulated and EGF-stimulated conditions **(Figure 3 a,b**), confirming the mass spectrometry results. As anticipated, treatment with MK-1175 reduced the hyperphosphorylation of CDK1 induced by Hybrid 7 (**Figure 3**).

**Figure 3.**
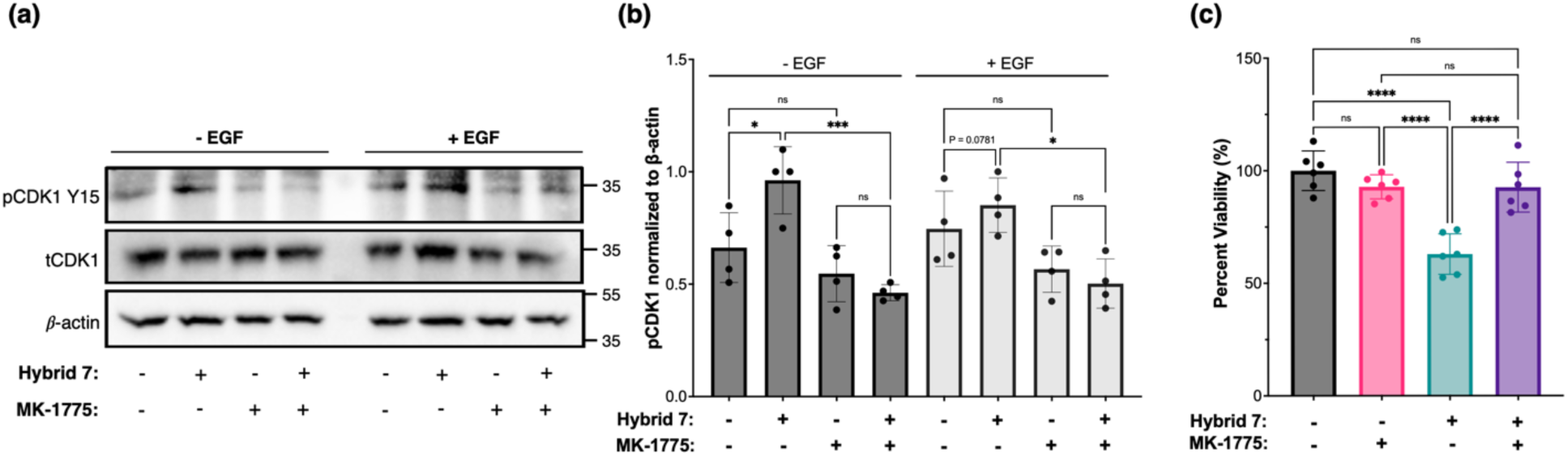
Increased levels of pCDK1 Y15 by Hybrid 7 treatment can be inhibited with a WEE-1 kinase inhibitor (MK-1775). A549 cells were treated with ± MK-1775 (200 nM) for 1 hour at 37°C, and then ± Hybrid 7 (10 µM) for 5 minutes at normal pH, pH adjusted to 5.5 for 10 min, and stimulated with 10 ng/mL EGF for 10 minutes before lysis. **(a)** Representative blots and quantification of phospho-protein levels of **(b)** CDK1 (Y15) are reported. Quantification of phospho-protein was normalized to β-actin control. **(c)** Following the same treatment with MK-1175 and Hybrid 7, cells were recovered for 72 h in the original media. Cell viability was assessed by MTT assay. All measurements are normalized to cells treated with no Hybrid 7 and no MK-1775 (100% viability). Results are shown as mean ± S.D., and the statistical significance of differences between samples was determined by one-way ANOVA with Tukey’s multiple comparisons correction (α = 0.05). ****, p ≤ 0.0001; ***, p ≤ 0.001; *, p ≤ 0.05; ns, p > 0.05. Only a subset of comparisons is shown for clarity.

To validate that CDK1 hyperphosphorylation plays a role in the anti-proliferative effect of Hybrid 7, an MTT assay was conducted with MK-1775 added prior to Hybrid 7 treatment. After treatment and a 72-hour recovery period, cells treated with Hybrid 7 alone (10 µM) exhibited a characteristic loss of proliferation compared to untreated control cells (**Fig. 3c**). As anticipated, treatment with MK-1775 alone did not significantly affect cell proliferation. However, pretreatment with MK-1775 before Hybrid 7 significantly restored proliferation compared to cells treated only with Hybrid 7, supporting the idea that the reduced proliferation is due to CDK1 hyperphosphorylation. Furthermore, no significant differences in proliferation were observed between cells treated with MK-1775 plus Hybrid 7 and those treated with MK-1775 alone or the untreated control, indicating that MK-1775 effectively restored proliferation to control-like levels. Altogether, results from immunoblotting and MTT assays support our hypothesis that CDK1 hyperphosphorylation is central to the inhibition of cell proliferation and migration observed after Hybrid 7 treatment. Having linked CDK1 hyperphosphorylation to reduced proliferation, we evaluated whether Hybrid 7 elevates DNA damage and oxidative stress in treated cells.

### Hybrid 7 increases DNA damage and intracellular ROS

To assess DNA damage, we quantified phosphorylation of Histone H2AX (γH2AX) at Serine 139 (pS139), a key marker for DNA damage that specifically indicates the presence of double-strand breaks (DSBs).^80–82^ After treatment with Hybrid 7 in EGF-stimulated or unstimulated conditions, cells were imaged and quantified for the intensity of γH2AX (pS139) localized within the nuclei (**Fig. 4**). Upon Hybrid 7 treatment, the signal significantly increased in the unstimulated and stimulated conditions (about a 1.7-fold increase in both cases; **Fig. 4b**). Another common feature of DNA damage is the formation of distinct foci due to the accumulation of phosphorylated γH2AX to repair DSBs.^81–85^ Foci are apparent in both samples treated with Hybrid 7. Because oxidative stress can promote double-strand breaks and checkpoint activation, we quantified the production of reactive oxygen species (ROS) with CellROX Green. A 10-min Hybrid 7 (10 µM) treatment, followed by a 1-h recovery, increased nuclear CellROX fluorescence relative to untreated cells, whereas low-pH buffer alone caused a smaller rise (**Fig. 4c and Fig. S3**). Hybrid 7 yielded ∼2-fold higher fluorescence than low pH alone, indicating ROS induction beyond acidification. While these measurements do not establish causality, they reveal a redox phenotype that coincides with the increase in γH2AX described above.

**Figure 4.**
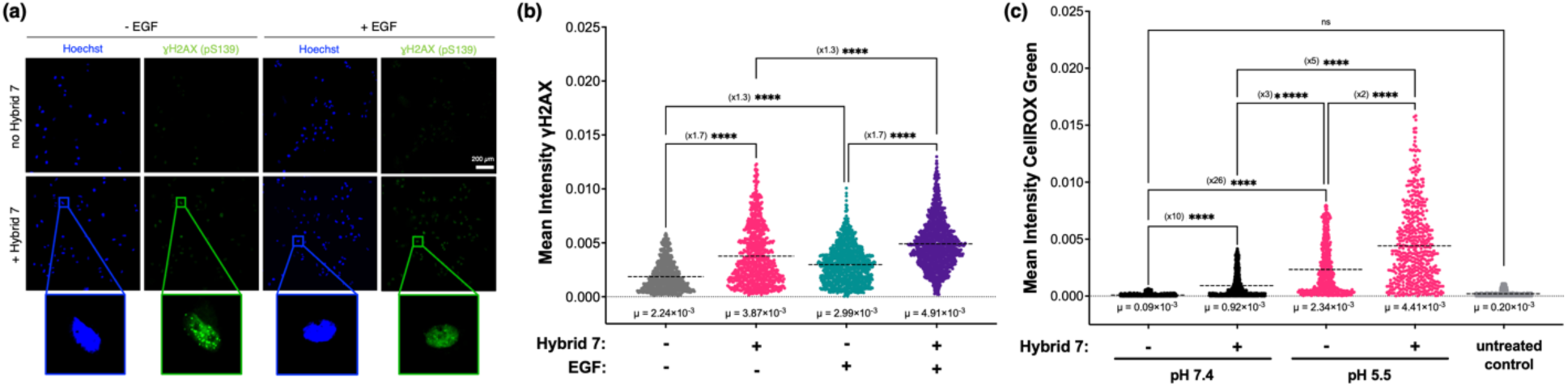
Hybrid 7 causes an increase in γH2AX phosphorylation and ROS production. **(a)** Representative immunostaining fluorescent images and **(b)** quantification of γH2AX phosphorylation after Hybrid 7 treatment (10 µM; 10 min) and one-hour incubation in the absence or presence of EGF stimulation. **(c)** Quantification of reactive oxygen species production using the membrane-permeable CellROX Green reagent after 10-minute Hybrid 7 treatment (10 µM). Cells were recovered in media for 1 hour before adding the CellROX stain. Results are shown as mean ± S.D. (n=3). Outlier points were identified using the ROUT method (Q = 1%) and removed. The statistical significance of differences between samples was determined by one-way ANOVA with Tukey’s multiple comparisons correction (α = 0.05). ****, p ≤ 0.0001; ***, p ≤ 0.001; *, p ≤ 0.05; ns, p > 0.05. Only a subset of comparisons is shown for clarity.

### Hybrid 7 induces G2/M arrest

Given the accumulation of DNA damage, increase in ROS production, and involvement of checkpoint signaling (manifested as inhibitory phosphorylation of CDK1) observed after Hybrid 7 treatment, we next asked whether these perturbations induce cell-cycle arrest. We quantified cell cycle distribution by propidium iodide staining and flow cytometry. Treatment with Hybrid 7 produced a ∼2-fold increase in the G2/M population with a corresponding decrease in G0/G1 under both EGF-stimulated and unstimulated conditions, consistent with G2/M arrest (**Fig. 5**). Together with the hyperphosphorylation of CDK1 at Y15, these results are consistent with checkpoint engagement after Hybrid 7 treatment.

**Figure 5.**
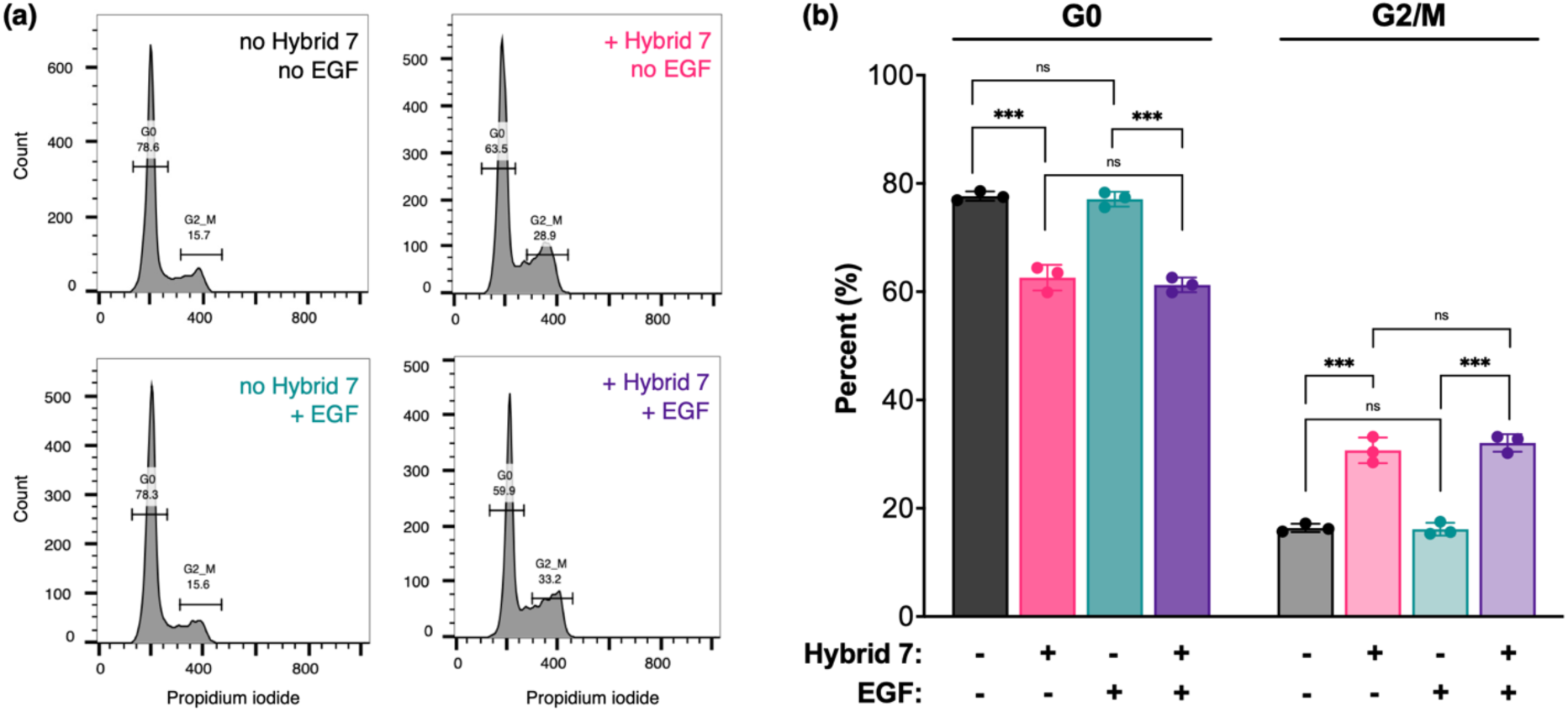
Hybrid 7 induces G2/M cell cycle arrest. A549 cells were treated with Hybrid 7 (10 µM, 10 minutes) followed by a 10-minute EGF stimulation (10 ng/mL) and recovered for 18 hours before staining with propidium iodide for cell cycle analysis. (**a)** Representative DNA content histograms with approximate G0 and G2/M population gating with **(b)** quantification of each cell cycle phase, indicating a significant decrease in G0 and an increase in G2/M for samples treated with Hybrid 7, regardless of growth factor background. Results are shown as mean ± S.D. (n=3). Statistical significance was determined by a two-tailed *t*-test. ***, p ≤ 0.001; ns, p > 0.05. Only a subset of comparisons is shown for clarity.

## Discussion

This study demonstrates that treatment with a pH-targeted transmembrane peptide agonist of PTPRJ (Hybrid 7) is sufficient to alter signaling and stress-response networks in A549 lung cancer cells and suppress proliferation and migration. Mechanistically, Hybrid 7 decreases EGFR phosphorylation, reduces phosphorylation across additional RTKs and motility-associated adaptors, elevates γH2AX and ROS, and drives inhibitory phosphorylation of CDK1 at Y15 with a corresponding G2/M accumulation. Pharmacological relief of that checkpoint with the WEE1 inhibitor MK-1775 restores proliferation, functionally linking CDK1 hyperphosphorylation to the cytostatic phenotype. Importantly, these effects occur in the genetic context of A549 (oncogenic *KRAS*^G12S^, *STK11*^Q37*^, *EGFR*^wt^, and *TP53*^wt^)^86–88^, which helps explain both the pattern and magnitude of the observed responses. We also note that Hybrid 7’s effects appear larger under unstimulated conditions than after acute EGF stimulation. This likely reflects ligand-driven saturation of kinase input (masking proportional dephosphorylation at a single time point) and EGFR internalization that reduces PTPRJ access to receptor sites.^89^

### PTPRJ agonism attenuates EGFR-centered signaling in A549

A549 cells are a non-EGFR-addicted lung adenocarcinoma model with wild-type EGFR and relatively modest receptor abundance. Yet, they maintain an EGFR pathway capable of stimulating migration and survival cues upon EGF exposure. However, constitutive KRAS activation due to the *KRAS*^G12S^ mutation uncouples major mitogenic outputs from receptor input.^90,91^ In this signaling and genetic context, attenuating EGFR phosphorylation by enhancing PTPRJ activity would be expected to blunt motility more readily than bulk mitogenic output. Consistent with this, Hybrid 7 decreased EGFR phosphorylation and reduced EGF-dependent migration, while proliferative signaling downstream of KRAS (e.g., Akt/ERK) remained largely intact at the sampled time point. The broader panel of Hybrid 7-sensitive sites identified by phosphoproteomics also underscores that enhancing a tumor-suppressive phosphatase at the membrane yields distributed, site-specific dephosphorylation across multiple RTKs and adaptors rather than a single-node effect. Importantly, proteins such as Src/Lyn family kinases, PI3K-p85, EphA2, and AXL collectively feed cytoskeletal remodeling, adhesion turnover, and directional motility.^92–97^ Such broad but low-amplitude changes could blunt motility and survival scaffolds while leaving some downstream averages apparently unchanged.

### From receptor dephosphorylation to checkpoint enforcement

Beyond RTKs, the most striking phosphoproteomic change was a robust increase in CDK1 Y15 phosphorylation under basal and EGF-stimulated conditions. Phosphorylation of Y15 is the canonical WEE1-installed inhibitory mark that enforces the G2/M checkpoint downstream of ATR/CHK1 or ATM/CHK2 upon DNA damage or replication stress.^98^ Consistent with activation of this axis, Hybrid 7 increased nuclear γH2AX and shifted cell-cycle distribution toward G2/M, and WEE1 inhibition with MK-1775 reduced CDK1-Y15 and rescued proliferation, indicating that Hybrid 7 primarily imposes a cytostatic checkpoint rather than lethal damage at the tested dose and schedule.

Together, these data also position CDK1 inhibition as an important effector of the cytostatic response to PTPRJ agonism in A549.

### ROS as a mechanistic bridge between receptor signaling, CDK1, and DNA damage

Short treatment with Hybrid 7 at low pH increased nuclear CellROX fluorescence beyond the effect of acidification alone, indicating a peptide-dependent redox disturbance sufficient to generate DNA lesions and stabilize γH2AX foci. Two non-exclusive mechanisms may link PTPRJ-driven receptor dephosphorylation to checkpoint engagement. First, by dampening EGFR activity and other RTKs, Hybrid 7 may reduce pro-survival and pro-repair inputs (e.g., PI3K/AKT, SFKs, DNA-dependent protein kinase), lowering the threshold for DDR activation in response to endogenous lesions. Second, EGFR and several other RTKs regulated by PTPRJ activity (e.g., PDGFβR, VEGFR2, MET, and FLT3) are linked to modulation of NADPH oxidases (NOX).^99–102^ Receptor dephosphorylation can acutely disrupt NOX regulation, causing transient alterations in ROS production. Once the DDR is initiated, it can reinforce itself by increasing cellular ROS, thereby maintaining checkpoint signaling and WEE1-dependent CDK1 Y15 inhibition until damage is resolved.^103^ This working model also provides a mechanistic basis for the strong anti-migratory effect observed under EGF stimulation and for inhibition of proliferation even without EGF, the latter arising from DDR checkpoint activation. This redox/checkpoint link is also compatible with A549’s genotype. *STK11* loss compromises AMPK-dependent energy and antioxidant programs, predisposing cells to oxidative stress and a heavier reliance on downstream checkpoints once damage accrues.^88,104^ KRAS activation dampens canonical mitogenic outputs, channeling the dominant phenotype toward motility suppression and checkpoint-driven cytostasis rather than acute apoptosis.^105^ Wild-type TP53 preserves damage sensing and checkpoint integrity, enabling durable G2/M arrest upon Hybrid 7 treatment.^87^ Experimental validation in a larger, genetically diverse panel of cell lines (KRAS/STK11/TP53/EGFR, variable PTPRJ and RTK expression) is underway but lies beyond the scope of this study.

## Conclusion

Activating PTPRJ with a pH-targeted transmembrane peptide in A549 cells leads to a two-pronged effect: receptor-proximal dephosphorylation that disrupts EGFR-linked motility networks, and a ROS→DDR→WEE1–CDK1 axis that enforces G2/M arrest and suppresses proliferation. The fact that even a short treatment (10 min) with a tumor-pH–guided peptide can induce lasting phenotypic effects highlights the potential of RPTP-targeting peptides as both tools to study RPTP biology and cancer therapeutics, alone or in combination (e.g., RTK inhibitors or radiotherapy to magnify redox stress).

## Materials and Methods

### Solid-phase peptide synthesis

Hybrid peptides were prepared by solid-phase peptide synthesis, purified and analyzed by RP-HPLC, and confirmed via MALDI-TOF mass spectrometry, as previously reported.^29,106^

### Cell Culture

Human lung carcinoma epithelial A549 cells and the human squamous cell carcinoma HSC3 cells were cultured in DMEM high glucose media supplemented with 10% fetal bovine serum (FBS), 100 units/mL penicillin, and 0.1 mg/mL streptomycin. Engineered human tongue squamous cell carcinoma HSC3 cells previously engineered to express wild-type PTPRJ ectopically (Bloch et al., 2019) were cultured in DMEM high glucose supplemented with 10% FBS, 100 units/mL penicillin, and 0.1 mg/mL streptomycin, and 2 µg/mL puromycin.^29^ All cell lines are grown and maintained in a humidified atmosphere of 5 % CO_2_ incubator at 37°C.

### A549 Treatment with Hybrid 7 and Assessment of Phosphorylation Levels

A549 cells were seeded in a 12-well plate to reach near confluency (∼90%). Cells were starved with DMEM supplemented with 1% FBS for 18 hours prior to treatment. Low pH media (pH 2.0, buffered with 25 mM sodium citrate and titrated with concentrated HCl) were made using media from an 18-hour starvation. The peptide is resuspended in DMSO (final concentration in solution does not exceed 0.5%) and sonicated briefly to facilitate solubilization. Cells were treated with 10 µM of peptide (or DMSO control) for 5 minutes at 37°C, 5% CO_2_. The cells were then titrated with pH 2 media to the desired pH of treatment (pH of ∼ 5.5) and incubated for 10 minutes at 37°C, 5% CO_2_. Do a series of gentle, incomplete washes with pH 7.4 media and prevent exposing cells to air. 10 ng/mL of EGF (Peprotech) was added for a 10-minute treatment at 37°C, 5% CO_2._ Immediately after, the media is aspirated and washed in cold 1X phosphate-buffered saline (PBS). Cells were lysed with cell extraction buffer supplemented with broad-spectrum phosphatase and protease inhibitors (Pierce #88667), spun down, and the supernatant was mixed with loading buffer. For MK-1775 treatment, after an 18-hour starvation, cells were treated with 200 µM MK-1775 (or DMSO control) for 1 hour at 37°C. The cells were then treated with 10 µM Hybrid for 5 minutes, pH adjusted to 5.5 for 10 min and stimulated with 10 ng/mL EGF for 10 minutes before lysis as previously described.

Samples were boiled for 10 minutes at 95°C and resolved by SDS-PAGE on a 10% tris-glycine gel. Subsequently, samples were transferred onto a 0.45 µM nitrocellulose membrane (GE Healthcare #1060002) at 25 Amp for 30 minutes using the TurboBlot system (Bio-Rad). Membranes were blocked in 5% w/v BSA in 1X tris-buffered saline Tween-20 (TBS-T) for 1 hour at room temperature and blotted for desired targets: phospho-EGF Receptor Tyr1068 (Cell Signaling Technology (CST) #3777), phospho-EGF Receptor Tyr1086 (CST #2220), phospho-cdc2 (CDK1) Tyr15 (CST #4539), phospho-histone H2A.X Ser139 (CST #9718), total EGFR (CST #4267), total cdc2 (CDK1) (CST, #9116) β-actin (CST #3700), and total PTPRJ (Santa Cruz #376794). The membranes were incubated with the appropriate antibody diluted in 5% TBS overnight (18h) at 4°C (pEGFR and tEGFR 1:1000 dilution, β-actin 1:4000 dilution, tPTPRJ 1:200 dilution). After successive washes, membranes were incubated with appropriate secondary antibodies at a 1:4000 dilution in TBS for 30 minutes at room temperature (anti-rabbit HRP-conjugated: CST #7074; anti-mouse HRP-conjugated: CST #7076). Following washes in TBS-T, membranes were visualized by chemiluminescence after adding Clarity Western ECL substrate (Bio-Rad #1705062). Images were quantified using ImageJ and plotted as normalized mean values (ratio of phosphorylated to total protein intensity).

### Wound Healing Scratch Assay

A549 cells were seeded into a 12-well plate at a cell density sufficient to achieve near confluency after 2 days, resulting in a nearly confluent monolayer (∼90%). 18 hours before treatment, cells were starved in DMEM containing 1% FBS. Peptide preparation and cell treatment were performed as described above. However, immediately after the 10-minute pH treatment, the monolayer of cells is scratched with a 200 µL pipette tip, and the debris is removed by the following incomplete washes as detailed above. The cells are then stimulated with 10 ng/mL EGF, which remains in the media for the duration of the wound closure (24 hours). Scratch areas were quantified using the MRI Wound Healing Tool in ImageJ, and the closure was determined by calculating the percent change in the area between the initial and final time points, reported as a percent closure value (%).

### MTT Proliferation Assay

Approximately 5,000 A549 cells were plated per well in a 96-well plate and allowed to adhere overnight. The cells were successively treated with serial dilutions of Hybrid 7 (starting at 10 µM) from a concentrated DMSO stock, as previously described. Following peptide treatment, the cells were resuspended in the original media from seeding to a final volume of 100 µL, and allowed to recover for 72h at 37°C. For MTT assays containing CDK1 inhibitor (MK-1775), after the cells were seeded and allowed to adhere overnight, they were treated with 200 nM MK-1775 (concentrated stock in DMSO) for 1 hour at 37°C, and then successively treated with Hybrid 7 as previously described. The cells were recovered in initial media for 72 hours at 37°C before analysis and readout. After a recovery period, MTT reagent (5 mg/mL stock in 1x PBS, 10 µL per 100 µL of media in each well) was added to each well and incubated for 3 hours at 37°C. The media was successively aspirated, and the remaining crystals were solubilized in DMSO before readout. Absorbance at 580 nm was measured, and the cell viability was calculated and normalized to the cells treated with pH 7.4 media only (100% viability). Data were fitted as a sigmoidal dose-response curve, and the IC_50_ value was determined via Prism 9 (GraphPad).

### Immunofluorescence Microscopy

To measure changes in Ki-67, A549 cells were seeded on 22 mm × 22 mm coverslips to achieve a confluence of ∼ 40-50% after 16 hours. Cells were treated with Hybrid 7 (10 µM for 10 min at pH 5.5), stimulated with 10 ng/mL EGF, and then incubated for 18 hours. After treatment, cells were washed with PBS and fixed with 4% paraformaldehyde in PBS for 20 minutes at room temperature.

Following additional PBS washes, cells were then permeabilized with 0.25% Triton X-100 in PBS for 5 minutes at room temperature. After another wash with PBS, the primary anti-Ki-67 mouse antibody (CST #9449) diluted in Intercept Blocking Buffer (Licor # 927-70001) was added, and the coverslips were incubated in a humidifying chamber for 18 hours at 4°C. After successive washes in PBS-Tween 20 (PBS-T), coverslips were incubated with anti-mouse Alexa Fluor 488 (Invitrogen #A11008) diluted (1:750) in Intercept Blocking Buffer for 1 hour at 37°C in a humidifying chamber. Following the final washes in PBS-T, coverslips were mounted onto slides with the Fluoromount mounting media (Sigma-Aldrich #F4680) and allowed to dry. Slides were imaged with a Nikon Eclipse Ti microscope with a 20x objective.

Immunofluorescence microscopy for DNA damage marker γH2AX was conducted using a similar workflow. Cells were seeded on coverslips, allowed to adhere overnight (∼60-70% confluent), and treated with Hybrid 7 as previously described. Cells were stimulated with 10 ng/mL EGF and allowed to recover for 1 hour at 37°C. Cells were fixed and permeabilized as described above. Coverslips were treated with a 1:400 dilution of primary anti-pSer 139 γH2AX antibody (CST #9718) in blocking buffer overnight at 4°C. Washes were conducted as previously described. The solution containing the secondary anti-rabbit Texas Red antibody (Invitrogen #T2767) was diluted (1:750) in blocking buffer, and the Hoechst nuclear DNA stain (Invitrogen #H3570) was added to coverslips and incubated at 37°C for 1 hour in a humidifying chamber. After the final washes, coverslips were mounted on slides and imaged as previously described. Smaller representative images were taken with the same objective (20x) using a smaller scanning area surrounding the cells (and nuclei) of interest. Images were processed using CellProfiler, and the intensities of Ki67 and nuclear H2AX were measured, quantified, and reported as mean intensity values. Representative images were processed via ImageJ.

### Multiplex Analysis

A549 cells were serum-starved in DMEM with 1% FBS overnight before Hybrid 7 treatment. Cells were treated with 10 µM Hybrid 7 for 5 minutes at 37°C, and pH was adjusted to pH 5.5 for 10 minutes at 37°C. After successive washes, cells were stimulated with EGF (10 ng/mL) for 10 minutes, followed by lysis in Cell Extraction Buffer (Thermo Fisher Scientific) supplemented with protease and phosphatase inhibitors. Total protein concentrations were determined using Pierce™ BCA Protein Assay Kits (Thermo Fisher Scientific). Luminex kits (Millipore Sigma) were used to measure relative phosphoprotein abundance according to the manufacturer’s instructions. Five or ten μg of total protein was loaded per well for the 8-plex MAPK/SAPK and 11-plex Akt/mTOR kits, respectively. Measurements were gathered using a Luminex xMAP Intelliflex system in the University of Virginia Flow Cytometry Core Facility.

### Mass Spectrometry Phosphoproteomics

<u>Cell treatment, lysis, and trypsinization</u>: A549 cells were treated with Hybrid 7 and stimulated with EGF for 10 minutes, as described above. Cells were lysed in 8 M urea lysis buffer, and the lysates were frozen at −80°C until use. After centrifugation at 4,200 × *g* for 10 min at 4°C, protein concentrations of the supernatant were measured by BCA. For each sample, 1 mg protein was diluted to 1 mg/mL with 8 M urea and was reduced with 10 mM dithiothreitol for 1 hour at 56°C, followed by alkylation with 55 mM iodoacetamide for 1 hour at room temperature, protected from light. The lysates were then diluted with 8 mL of 100 mM ammonium acetate, pH 8.9, and digested with sequencing-grade trypsin (Promega) at a trypsin-to-protein ratio of 1:50, overnight at room temperature. The trypsinization reaction was quenched by adding 1 mL acetic acid, and the peptides were cleaned up with Sep-Pak C18 cartridges (Waters). Peptides were eluted with 7 mL 40% acetonitrile in 0.1% acetic acid. The elution was concentrated, and acetonitrile was removed by SpeedVac vacuum concentrators (Thermo Fisher Scientific). After measuring concentrations by BCA, 190 μg peptide aliquots from each sample were lyophilized.

<u>Tandem mass tag (TMT) labeling</u>: For tandem mass tag (TMT) labeling, 190 μg tryptic peptide aliquots from each sample were resuspended in 50 μL 50 mM HEPES, pH 8.5. 12 channels from a TMT 18-plex (Thermo Fisher Scientific) were chosen for labeling. 0.5 mg TMT reagent from each channel was resuspended in 15 μL of anhydrous acetonitrile and was added to the resuspended peptides. After incubation for 4 hours at room temperature on a shaker at 400 rpm, the labeling reactions were quenched by adding 3.2 μL 5% hydroxylamine. The labeling reactions were pooled, and the labeling tubes were washed twice with 40 μL of 25% acetonitrile in 0.1% acetic acid. The washes were then pooled together with the labeling reactions. The pooled solution was dried by SpeedVac and stored at −80°C until use.

<u>Phosphotyrosine (pY) enrichment</u>: Phosphotyrosine immunoaffinity purification (IP) was first applied to enrich tyrosine phosphorylated peptides. 60 μL protein G agarose bead slurry (Millipore) was conjugated to 24 μg 4G10 pY antibody (BioXCell) and 6 μL PT66 pY antibody (Sigma) for 5 hours at 4°C. The TMT-labeled dried peptides were resuspended in 400 μL IP buffer (100 mM Tris-HCl, 1% NP-40, pH 7.4), and mixed with the conjugated beads and incubated overnight on a rotator at 4°C. Beads were then washed once with the IP buffer, and three times with rinse buffer (100 mM Tris-HCl, pH 7.4). Tyrosine-phosphorylated peptides were eluted twice with 25 μL 0.2% trifluoroacetic acid (TFA) at room temperature for 10 min each. The IP elution was cleaned up, and the phosphopeptides were further enriched by High-Select™ ferric nitrilotriacetate (Fe-NTA) Phosphopeptide Enrichment Kit (Thermo Fisher Scientific) according to the manufacturer’s protocol. The final phosphopeptide elution was dried down by SpeedVac, resuspended in 5 μL 3% acetonitrile in 0.1% formic acid, and was ready for LC/MS-MS analysis. The supernatant of pY IP (which contains total peptides other than pY peptides) was diluted 500-fold with 0.1% formic acid, and 3 μL of the diluted supernatant was used for a crude total peptide MS analysis to normalize the peptide loadings of each TMT channel.

<u>Liquid Chromatography with tandem mass spectrometry (LC/MS-MS) analysis</u>: The phosphopeptide MS sample was loaded, using a column-packing device, onto a homemade laser-pulled analytical capillary column with 50 μm inner diameter and 10 cm length (Polymicro), manually packed with 3 μm C18 beads (YMC-Gel). The MS experiment was performed with an Orbitrap Exploris 480 mass spectrometer (Thermo Fisher Scientific) coupled to an Agilent 1100 Series HPLC. Peptides were separated with the following 140 min gradient consisting of 0.2 M acetic acid (buffer A) and 70% acetonitrile in 0.2 M acetic acid (buffer B): 0 – 10 min: 0% to 11% B, 10 – 115 min: 11% to 32% B, 115 – 125 min: 32% to 60% B, 125-130 min: 60% to 100% B, 130 – 133 min: 100% B, 133 – 140 min: 100% to 0% B. The LC was operated at a flow rate of 0.2 mL/min to ensure a nanoliter-level flow rate through the analytical column. The MS run was done in data-dependent acquisition (DDA) mode, with the following parameters for MS1: Orbitrap resolution 120000; m/z scan range 380-1800; normalized AGC target 300%; maximum injection time (IT) 50 ms; only precursors with charge states 2 to 6 were included. For dynamic exclusion, ions occurred 3 times within 30 s were excluded for 120 s, and the exclusion window was +/- 3 ppm. The MS2 parameters are as follows: isolation window 0.4 m/z, normalized HCD collision energy (NCE) 33%, orbitrap resolution 120000, normalized AGC target 1000%, maximum infection time 247 ms.

The crude total peptide sample was bomb loaded onto a homemade precolumn with 100 μm inner diameter and 10 cm length (Polymicro), packed with 10 μm C18 beads (YMC-Gel), and the precolumn was connected to an analytical column. The MS analysis was done on Q Exactive Plus (Thermo Fisher Scientific), with a 60 min linear gradient of the same buffer A and B. The MS run was done in DDA mode, with the following MS1 parameters: resolution 70000; AGC target 3 x 10^6^; maximum IT 50 ms; scan range 350-2000 m/z. The top 10 ions were fragmented for MS2 analysis with following parameters: resolution 35000; AGC target 1 x 10^5^; maximum IT 150 ms; isolation window 0.4 m/z; NCE 33%; dynamic exclusion 30s.

<u>Peptide identification and quantification</u>: Mass spectra raw files were searched by Proteome Discoverer 3.0 (Thermo Fisher Scientific). The search was conducted by Mascot search engine (Matrix Science) against the human SwissProt database. The search allowed a maximum of 2 miss cleavage sites, and the mass tolerances were 20 mmu for fragments and 10 ppm for precursors. TMT-labeled lysine, TMT-labeled peptide N-terminus, and cysteine carbamidomethylation were searched as static modifications. Serine, threonine, and tyrosine phosphorylation, as well as methionine oxidation were searched as dynamic modifications. Peptide spectrum matches (PSMs) with a minimum peptide length of 6, search engine rank = 1, and mascot ion score (IS) > 20 were kept for further analyses. PSMs with missing values across any channel were filtered out. The data analyses were conducted with RStudio and Microsoft Excel. TMT reporter ion intensities from PSMs were summed for each unique phosphopeptide. For each channel, the peptide abundances were normalized to the total intensities of the crude total peptide analysis, to correct the sample loading variances of TMT channels. A student t-test was applied to determine the statistical significance of different conditions.

## Data availability

The mass spectrometry data have been deposited to the ProteomeXchange Consortium via the PRIDE partner repository with the dataset identifier PXD061999.^107–109^

Reviewers can access the dataset by logging in to the PRIDE website using the following account details: Username; Password: Lhskdi1m9OIj

### Cell Cycle Analysis

A549 cells were seeded in 6-well plates to achieve ∼60% confluence by the next day. Cells were then starved overnight in 1% FBS, treated with Hybrid 7 (10 µM), and/or successively stimulated with 10 ng/mL EGF (left on for the duration of the incubation), as previously described. After the desired recovery time (18 hours), the cells were removed from the bottom of the wells with StemPro^TM^ Accutase^TM^ Cell Dissociation reagent (Gibco, #A1110501), spun down, and successively washed. The cells were resuspended in 1X PBS, and ice-cold 100% ethanol was added dropwise to the cells while vortexing to prevent clumping, resulting in a final concentration of 70% ethanol. The cells were stored at −20°C before staining. Fixed cells were washed and stained with 50 µg/mL PI supplemented with 100 µg/mL RNase A (CITE)in PBS for 30 minutes before analysis by flow cytometer.

The cells were analyzed using a BDFacs Canto II flow cytometer equipped with a 488 nm argon laser with a 585/20 bandpass filter. A minimum of 20,000 events was counted for each experimental condition. For analysis, cells were gated for the live cell population, and recorded only singlet measurements by excluding doublets via forward scattering thresholds. Cell Cycle Analysis was performed with FlowJo and by gating for the G0 and G2/M peaks of the total populations. Each cell cycle phase was reported as a percentage (%), respectively.

### Detection of Reactive Oxygen Species

A549 cells were seeded on 22 mm × 22 mm coverslips in 6-well plates at 200,000 cells per well to be ∼ 40-50% confluent after 16 hours. Cells were treated with the Hybrid 7 peptide as previously described at both high (pH 8) and low (pH 5.5) pH conditions, without subsequent EGF treatment. The cells were recovered in the initial media for 1 hour at 37°C. CellROX Green live cell stain for oxidized species (Invitrogen #C10444) (5 µM) was added to cells for 30 minutes at 37°C. Cells were washed, fixed, and stained with a DNA nuclear stain as previously described above and mounted onto slides. Images were processed using CellProfiler, where the fraction positive measures each “positive” nucleus containing a mean CellROX signal over a threshold value (median intensity ± 2 standard deviations). The values reported are a ratio of “positive” nuclei measured to the total nuclear population.

## Acknowledgments

This work was supported by the National Institutes of Health under awards NIGMS R01 GM139998 (to D.T. and M.J.L), NCI P30 CA044579 (UVA Cancer Center Core), and U54 CA283114 (to F.M.W).

## Notes

### Competing Interest Statement

The authors have declared no competing interest.

## References

(1) Manning, G.; Whyte, D. B.; Martinez, R.; Hunter, T.; Sudarsanam, S. The Protein Kinase Complement of the Human Genome. Science 2002, 298 (5600), 1912–1934. 10.1126/science.1075762.

(2) Ubersax, J. A.; Ferrell Jr, J. E. Mechanisms of Specificity in Protein Phosphorylation. Nat Rev Mol Cell Biol 2007, 8 (7), 530–541. 10.1038/nrm2203.

(3) Day, E. K.; Sosale, N. G.; Lazzara, M. J. Cell Signaling Regulation by Protein Phosphorylation: A Multivariate, Heterogeneous, and Context-Dependent Process. Curr Opin Biotechnol 2016, 40, 185–192. 10.1016/j.copbio.2016.06.005.

(4) Tonks, N. K. Protein Tyrosine Phosphatases: From Genes, to Function, to Disease. Nat Rev Mol Cell Biol 2006, 7 (11), 833–846. 10.1038/nrm2039.

(5) Hunter, T. Tyrosine Phosphorylation: Thirty Years and Counting. Current Opinion in Cell Biology 2009, 21 (2), 140–146. 10.1016/j.ceb.2009.01.028.

(6) Tonks, N. K. Protein Tyrosine Phosphatases - from Housekeeping Enzymes to Master Regulators of Signal Transduction. FEBS Journal 2013, 280 (2), 346–378. 10.1111/febs.12077.

(7) Cohen, P.; Alessi, D. R. Kinase Drug Discovery – What’s Next in the Field? ACS Chem. Biol. 2013, 8 (1), 96–104. 10.1021/cb300610s.

(8) He, R.; Yu, Z.; Zhang, R.; Zhang, Z. Protein Tyrosine Phosphatases as Potential Therapeutic Targets. Acta Pharmacol Sin 2014, 35 (10), 1227–1246. 10.1038/aps.2014.80.

(9) Libermann, T. A.; Nusbaum, H. R.; Razon, N.; Kris, R.; Lax, I.; Soreq, H.; Whittle, N.; Waterfield, M. D.; Ullrich, A.; Schlessinger, J. Amplification, Enhanced Expression and Possible Rearrangement of EGF Receptor Gene in Primary Human Brain Tumours of Glial Origin. Nature 1985, 313 (5998), 144–147. 10.1038/313144a0.

(10) Mok, T. S. K. Personalized Medicine in Lung Cancer: What We Need to Know. Nat Rev Clin Oncol 2011, 8 (11), 661–668. 10.1038/nrclinonc.2011.126.

(11) Spano, J.-P.; Lagorce, C.; Atlan, D.; Milano, G.; Domont, J.; Benamouzig, R.; Attar, A.; Benichou, J.; Martin, A.; Morere, J.-F.; Raphael, M.; Penault-Llorca, F.; Breau, J.-L.; Fagard, R.; Khayat, D.; Wind, P. Impact of EGFR Expression on Colorectal Cancer Patient Prognosis and Survival. Annals of Oncology 2005, 16 (1), 102–108. 10.1093/annonc/mdi006.

(12) Ercan, D.; Xu, C.; Yanagita, M.; Monast, C. S.; Pratilas, C. A.; Montero, J.; Butaney, M.; Shimamura, T.; Sholl, L.; Ivanova, E. V.; Tadi, M.; Rogers, A.; Repellin, C.; Capelletti, M.; Maertens, O.; Goetz, E. M.; Letai, A.; Garraway, L. A.; Lazzara, M. J.; Rosen, N.; Gray, N. S.; Wong, K.-K.; Jänne, P. A. Reactivation of ERK Signaling Causes Resistance to EGFR Kinase Inhibitors. Cancer Discovery 2012, 2 (10), 934–947. 10.1158/2159-8290.CD-12-0103.

(13) Furcht, C. M.; Muñoz Rojas, A. R.; Nihalani, D.; Lazzara, M. J. Diminished Functional Role and Altered Localization of SHP2 in Non-Small Cell Lung Cancer Cells with EGFR-Activating Mutations. Oncogene 2013, 32 (18), 2346–2355. 10.1038/onc.2012.240.

(14) Lazzara, M. J.; Lane, K.; Chan, R.; Jasper, P. J.; Yaffe, M. B.; Sorger, P. K.; Jacks, T.; Neel, B. G.; Lauffenburger, D. A. Impaired SHP2-Mediated Extracellular Signal-Regulated Kinase Activation Contributes to Gefitinib Sensitivity of Lung Cancer Cells with Epidermal Growth Factor Receptor–Activating Mutations. Cancer Research 2010, 70 (9), 3843–3850. 10.1158/0008-5472.CAN-09-3421.

(15) Niederst, M. J.; Engelman, J. A. Bypass Mechanisms of Resistance to Receptor Tyrosine Kinase Inhibition in Lung Cancer. Sci. Signal. 2013, 6 (294). 10.1126/scisignal.2004652.

(16) Hata, A. N.; Niederst, M. J.; Archibald, H. L.; Gomez-Caraballo, M.; Siddiqui, F. M.; Mulvey, H. E.; Maruvka, Y. E.; Ji, F.; Bhang, H. C.; Krishnamurthy Radhakrishna, V.; Siravegna, G.; Hu, H.; Raoof, S.; Lockerman, E.; Kalsy, A.; Lee, D.; Keating, C. L.; Ruddy, D. A.; Damon, L. J.; Crystal, A. S.; Costa, C.; Piotrowska, Z.; Bardelli, A.; Iafrate, A. J.; Sadreyev, R. I.; Stegmeier, F.; Getz, G.; Sequist, L. V.; Faber, A. C.; Engelman, J. A. Tumor Cells Can Follow Distinct Evolutionary Paths to Become Resistant to Epidermal Growth Factor Receptor Inhibition. Nat Med 2016, 22 (3), 262–269. 10.1038/nm.4040.

(17) Nunes-Xavier, C. E.; Martín-Pérez, J.; Elson, A.; Pulido, R. Protein Tyrosine Phosphatases as Novel Targets in Breast Cancer Therapy. Biochimica et Biophysica Acta (BBA) - Reviews on Cancer 2013, 1836 (2), 211–226. 10.1016/j.bbcan.2013.06.001.

(18) Ruivenkamp, C. A. L.; van Wezel, T.; Zanon, C.; Stassen, A. P. M.; Vlcek, C.; Csikós, T.; Klous, A. M.; Tripodis, N.; Perrakis, A.; Boerrigter, L.; Groot, P. C.; Lindeman, J.; Mooi, W. J.; Meijjer, G. A.; Scholten, G.; Dauwerse, H.; Paces, V.; van Zandwijk, N.; van Ommen, G. J. B.; Demant, P. Ptprj Is a Candidate for the Mouse Colon-Cancer Susceptibility Locus Scc1 and Is Frequently Deleted in Human Cancers. Nat Genet 2002, 31 (3), 295–300. 10.1038/ng903.

(19) Iuliano, R.; Pera, I. L.; Cristofaro, C.; Baudi, F.; Arturi, F.; Pallante, P.; Martelli, M. L.; Trapasso, F.; Chiariotti, L.; Fusco, A. The Tyrosine Phosphatase PTPRJ/DEP-1 Genotype Affects Thyroid Carcinogenesis. Oncogene 2004, 23 (52), 8432–8438. 10.1038/sj.onc.1207766.

(20) Ruivenkamp, C.; Hermsen, M.; Postma, C.; Klous, A.; Baak, J.; Meijer, G.; Demant, P. LOH of PTPRJ Occurs Early in Colorectal Cancer and Is Associated with Chromosomal Loss of 18q12–21. Oncogene 2003, 22 (22), 3472–3474. 10.1038/sj.onc.1206246.

(21) Keane, M. M.; Lowrey, G. A.; Ettenberg, S. A.; Dayton, M. A.; Lipkowitz, S. The Protein Tyrosine Phosphatase DEP-1 Is Induced during Differentiation and Inhibits Growth of Breast Cancer Cells. Cancer Res 1996, 56 (18), 4236–4243.

(22) Farrington, C. C.; Yuan, E.; Mazhar, S.; Izadmehr, S.; Hurst, L.; Allen-Petersen, B. L.; Janghorban, M.; Chung, E.; Wolczanski, G.; Galsky, M.; Sears, R.; Sangodkar, J.; Narla, G. Protein Phosphatase 2A Activation as a Therapeutic Strategy for Managing MYC-Driven Cancers. Journal of Biological Chemistry 2020, 295 (3), 757–770. 10.1016/S0021-9258(17)49933-9.

(23) Takahashi, T.; Takahashi, K.; Mernaugh, R. L.; Tsuboi, N.; Liu, H.; Daniel, T. O. A Monoclonal Antibody against CD148, a Receptor-like Tyrosine Phosphatase, Inhibits Endothelial-Cell Growth and Angiogenesis. Blood 2006, 108 (4), 1234–1242. 10.1182/blood-2005-10-4296.

(24) Takahashi, K.; Kim, R. H.; Pasic, L.; He, L.; Nagasaka, S.; Katagiri, D.; May, T.; Shimizu, A.; Harris, R. C.; Mernaugh, R. L.; Takahashi, T. Agonistic Anti-CD148 Monoclonal Antibody Attenuates Diabetic Nephropathy in Mice. Am J Physiol Renal Physiol 2020, 318 (3), F647–F659. 10.1152/ajprenal.00288.2019.

(25) Bali, N.; Lee, H.-K. (Peter); Zinn, K. Sticks and Stones, a Conserved Cell Surface Ligand for the Type IIa RPTP Lar, Regulates Neural Circuit Wiring in Drosophila. eLife 2022, 11, e71469. 10.7554/eLife.71469.

(26) Tremblay, M. L. Advancing Therapeutics Using Antibody-Induced Dimerization of Receptor Tyrosine Phosphatases. Genes Dev 2023, 37 (15–16), 678–680. 10.1101/gad.351120.123.

(27) Stoker, A. W. Detection and Identification of Ligands for Mammalian RPTP Extracellular Domains. In *Protein Tyrosine Phosphatases*; Pulido, R., Ed.; Methods in Molecular Biology; Springer New York: New York, NY, 2016; Vol. 1447, pp 267–281. 10.1007/978-1-4939-3746-2_15.

(28) Jiang, G.; Den Hertog, J.; Su, J.; Noel, J.; Sap, J.; Hunter, T. Dimerization Inhibits the Activity of Receptor-like Protein-Tyrosine Phosphatase-α. Nature 1999, 401 (6753), 606–610. 10.1038/44170.

(29) Bloch, E.; Sikorski, E. L.; Pontoriero, D.; Day, E. K.; Berger, B. W.; Lazzara, M. J.; Thévenin, D. Disrupting the Transmembrane Domain–Mediated Oligomerization of Protein Tyrosine Phosphatase Receptor J Inhibits EGFR-Driven Cancer Cell Phenotypes. Journal of Biological Chemistry 2019, 294 (49), 18796–18806. 10.1074/jbc.RA119.010229.

(30) Tertoolen, L. G.; Blanchetot, C.; Jiang, G.; Overvoorde, J.; Gadella, T. W.; Hunter, T.; Hertog, J. den. Dimerization of Receptor Protein-Tyrosine Phosphatase Alpha in Living Cells. BMC Cell Biol 2001, 2 (1), 8. 10.1186/1471-2121-2-8.

(31) Chin, C.-N.; Sachs, J. N.; Engelman, D. M. Transmembrane Homodimerization of Receptor-like Protein Tyrosine Phosphatases. FEBS Letters 2005, 579 (17), 3855–3858. 10.1016/j.febslet.2005.05.071.

(32) Barr, A. J.; Ugochukwu, E.; Lee, W. H.; King, O. N. F.; Filippakopoulos, P.; Alfano, I.; Savitsky, P.; Burgess-Brown, N. A.; Müller, S.; Knapp, S. Large-Scale Structural Analysis of the Classical Human Protein Tyrosine Phosphatome. Cell 2009, 136 (2), 352–363. 10.1016/j.cell.2008.11.038.

(33) Hower, A. E.; Beltran, P. J.; Bixby, J. L. Dimerization of Tyrosine Phosphatase PTPRO Decreases Its Activity and Ability to Inactivate TrkC. Journal of Neurochemistry 2009, 110 (5), 1635–1647. 10.1111/j.1471-4159.2009.06261.x.

(34) Jandt, E.; Denner, K.; Kovalenko, M.; Östman, A.; Böhmer, F.-D. The Protein-Tyrosine Phosphatase DEP-1 Modulates Growth Factor-Stimulated Cell Migration and Cell–Matrix Adhesion. Oncogene 2003, 22 (27), 4175–4185. 10.1038/sj.onc.1206652.

(35) Kovalenko, M.; Denner, K.; Sandström, J.; Persson, C.; Groβ, S.; Jandt, E.; Vilella, R.; Böhmer, F.; Östman, A. Site-Selective Dephosphorylation of the Platelet-Derived Growth Factor β-Receptor by the Receptor-like Protein-Tyrosine Phosphatase DEP-1. Journal of Biological Chemistry 2000, 275 (21), 16219–16226. 10.1074/jbc.275.21.16219.

(36) Petermann, A.; Haase, D.; Wetzel, A.; Balavenkatraman, K. K.; Tenev, T.; Gührs, K.-H.; Friedrich, S.; Nakamura, M.; Mawrin, C.; Böhmer, F.-D. Loss of the Protein-Tyrosine Phosphatase DEP-1/PTPRJ Drives Meningioma Cell Motility: Regulation of Meningioma Cell-Motility by DEP-1. Brain Pathology 2011, 21 (4), 405–418. 10.1111/j.1750-3639.2010.00464.x.

(37) Lampugnani, M. G.; Zanetti, A.; Corada, M.; Takahashi, T.; Balconi, G.; Breviario, F.; Orsenigo, F.; Cattelino, A.; Kemler, R.; Daniel, T. O.; Dejana, E. Contact Inhibition of VEGF-Induced Proliferation Requires Vascular Endothelial Cadherin, β-Catenin, and the Phosphatase DEP-1/CD148. Journal of Cell Biology 2003, 161 (4), 793–804. 10.1083/jcb.200209019.

(38) Patel, J. P.; Gönen, M.; Figueroa, M. E.; Fernandez, H.; Sun, Z.; Racevskis, J.; Van Vlierberghe, P.; Dolgalev, I.; Thomas, S.; Aminova, O.; Huberman, K.; Cheng, J.; Viale, A.; Socci, N. D.; Heguy, A.; Cherry, A.; Vance, G.; Higgins, R. R.; Ketterling, R. P.; Gallagher, R. E.; Litzow, M.; van den Brink, M. R. M.; Lazarus, H. M.; Rowe, J. M.; Luger, S.; Ferrando, A.; Paietta, E.; Tallman, M. S.; Melnick, A.; Abdel-Wahab, O.; Levine, R. L. Prognostic Relevance of Integrated Genetic Profiling in Acute Myeloid Leukemia. N Engl J Med 2012, 366 (12), 1079–1089. 10.1056/NEJMoa1112304.

(39) Palka, H. L.; Park, M.; Tonks, N. K. Hepatocyte Growth Factor Receptor Tyrosine Kinase Met Is a Substrate of the Receptor Protein-Tyrosine Phosphatase DEP-1. Journal of Biological Chemistry 2003, 278 (8), 5728–5735. 10.1074/jbc.M210656200.

(40) Avraham, R.; Yarden, Y. Feedback Regulation of EGFR Signalling: Decision Making by Early and Delayed Loops. Nat Rev Mol Cell Biol 2011, 12 (2), 104–117. 10.1038/nrm3048.

(41) Tarcic, G.; Boguslavsky, S. K.; Wakim, J.; Kiuchi, T.; Liu, A.; Reinitz, F.; Nathanson, D.; Takahashi, T.; Mischel, P. S.; Ng, T.; Yarden, Y. An Unbiased Screen Identifies DEP-1 Tumor Suppressor as a Phosphatase Controlling EGFR Endocytosis. Current Biology 2009, 19 (21), 1788–1798. 10.1016/j.cub.2009.09.048.

(42) Arora, D.; Stopp, S.; Böhmer, S.-A.; Schons, J.; Godfrey, R.; Masson, K.; Razumovskaya, E.; Rönnstrand, L.; Tänzer, S.; Bauer, R.; Böhmer, F.-D.; Müller, J. P. Protein-Tyrosine Phosphatase DEP-1 Controls Receptor Tyrosine Kinase FLT3 Signaling. Journal of Biological Chemistry 2011, 286 (13), 10918–10929. 10.1074/jbc.M110.205021.

(43) Böhmer, S.-A.; Weibrecht, I.; Söderberg, O.; Böhmer, F.-D. Association of the Protein-Tyrosine Phosphatase DEP-1 with Its Substrate FLT3 Visualized by In Situ Proximity Ligation Assay. PLoS ONE 2013, 8 (5), e62871. 10.1371/journal.pone.0062871.

(44) Schwarz, M.; Rizzo, S.; Paz, W. E.; Kresinsky, A.; Thévenin, D.; Müller, J. P. Disrupting PTPRJ Transmembrane-Mediated Oligomerization Counteracts Oncogenic Receptor Tyrosine Kinase FLT3 ITD. Front. Oncol. 2022, 12, 1017947. 10.3389/fonc.2022.1017947.

(45) Kim, J.; Dang, C. V. Cancer’s Molecular Sweet Tooth and the Warburg Effect: Figure 1. Cancer Res 2006, 66 (18), 8927–8930. 10.1158/0008-5472.CAN-06-1501.

(46) Gatenby, R. A.; Gillies, R. J. Why Do Cancers Have High Aerobic Glycolysis? Nat Rev Cancer 2004, 4 (11), 891–899. 10.1038/nrc1478.

(47) Helmlinger, G.; Sckell, A.; Dellian, M.; Forbes, N. S.; Jain, R. K. Acid Production in Glycolysis-Impaired Tumors Provides New Insights into Tumor Metabolism. Clin Cancer Res 2002, 8 (4), 1284–1291.

(48) Seyfried, T. N.; Shelton, L. M. Cancer as a Metabolic Disease. Nutr Metab (Lond) 2010, 7 (1), 7. 10.1186/1743-7075-7-7.

(49) Hanahan, D.; Weinberg, R. A. Hallmarks of Cancer: The Next Generation. Cell 2011, 144 (5), 646–674. 10.1016/j.cell.2011.02.013.

(50) Vander Heiden, M. G.; Cantley, L. C.; Thompson, C. B. Understanding the Warburg Effect: The Metabolic Requirements of Cell Proliferation. Science 2009, 324 (5930), 1029–1033. 10.1126/science.1160809.

(51) Rizzo, S.; Sikorski, E.; Park, S.; Im, W.; Vasquez-Montes, V.; Ladokhin, A. S.; Thévenin, D. Promoting the Activity of a Receptor Tyrosine Phosphatase with a Novel PH -responsive Transmembrane Agonist Inhibits Cancer-associated Phenotypes. Protein Science 2023, 32 (9), e4742. 10.1002/pro.4742.

(52) Arafeh, R.; Shibue, T.; Dempster, J. M.; Hahn, W. C.; Vazquez, F. The Present and Future of the Cancer Dependency Map. Nat Rev Cancer 2025, 25 (1), 59–73. 10.1038/s41568-024-00763-x.

(53) Yamaoka, T.; Frey, M. R.; Dise, R. S.; Bernard, J. K.; Polk, D. B. Specific Epidermal Growth Factor Receptor Autophosphorylation Sites Promote Mouse Colon Epithelial Cell Chemotaxis and Restitution. Am J Physiol Gastrointest Liver Physiol 2011, 301 (2), G368–376. 10.1152/ajpgi.00327.2010.

(54) Gerritsen, J. S.; Faraguna, J. S.; Bonavia, R.; Furnari, F. B.; White, F. M. Predictive Data-Driven Modeling of C-Terminal Tyrosine Function in the EGFR Signaling Network. Life Sci. Alliance 2023, 6 (8), e202201466. 10.26508/lsa.202201466.

(55) Lefebvre, C.; Allan, A. L. Anti-Proliferative and Anti-Migratory Effects of EGFR and c-Met Tyrosine Kinase Inhibitors in Triple Negative Breast Cancer Cells. Precis Cancer Med 2021, 4, 2–2. 10.21037/pcm-20-62.

(56) Barnes, C. J.; Bagheri-Yarmand, R.; Mandal, M.; Yang, Z.; Clayman, G. L.; Hong, W. K.; Kumar, R. Suppression of Epidermal Growth Factor Receptor, Mitogen-Activated Protein Kinase, and Pak1 Pathways and Invasiveness of Human Cutaneous Squamous Cancer Cells by the Tyrosine Kinase Inhibitor ZD1839 (Iressa)1. Molecular Cancer Therapeutics 2003, 2 (4), 345–351.

(57) Boucher, I.; Kehasse, A.; Marcincin, M.; Rich, C.; Rahimi, N.; Trinkaus-Randall, V. Distinct Activation of Epidermal Growth Factor Receptor by UTP Contributes to Epithelial Cell Wound Repair. The American Journal of Pathology 2011, 178 (3), 1092–1105. 10.1016/j.ajpath.2010.11.060.

(58) Salazar-Cavazos, E.; Nitta, C. F.; Mitra, E. D.; Wilson, B. S.; Lidke, K. A.; Hlavacek, W. S.; Lidke, D. S. Multisite EGFR Phosphorylation Is Regulated by Adaptor Protein Abundances and Dimer Lifetimes. MBoC 2020, 31 (7), 695–708. 10.1091/mbc.E19-09-0548.

(59) Ngan, E.; Stoletov, K.; Smith, H. W.; Common, J.; Muller, W. J.; Lewis, J. D.; Siegel, P. M. LPP Is a Src Substrate Required for Invadopodia Formation and Efficient Breast Cancer Lung Metastasis. Nat Commun 2017, 8 (1), 15059. 10.1038/ncomms15059.

(60) Dodelet, V. C.; Pazzagli, C.; Zisch, A. H.; Hauser, C. A.; Pasquale, E. B. A Novel Signaling Intermediate, SHEP1, Directly Couples Eph Receptors to R-Ras and Rap1A. Journal of Biological Chemistry 1999, 274 (45), 31941–31946. 10.1074/jbc.274.45.31941.

(61) Lu, Y.; Brush, J.; Stewart, T. A. NSP1 Defines a Novel Family of Adaptor Proteins Linking Integrin and Tyrosine Kinase Receptors to the C-Jun N-Terminal Kinase/Stress-Activated Protein Kinase Signaling Pathway. Journal of Biological Chemistry 1999, 274 (15), 10047–10052. 10.1074/jbc.274.15.10047.

(62) Jones, R. B.; Gordus, A.; Krall, J. A.; MacBeath, G. A Quantitative Protein Interaction Network for the ErbB Receptors Using Protein Microarrays. Nature 2006, 439 (7073), 168–174. 10.1038/nature04177.

(63) Bos, J. L.; De Rooij, J.; Reedquist, K. A. Rap1 Signalling: Adhering to New Models. Nat Rev Mol Cell Biol 2001, 2 (5), 369–377. 10.1038/35073073.

(64) Sakakibara, A.; Ohba, Y.; Kurokawa, K.; Matsuda, M.; Hattori, S. Novel Function of Chat in Controlling Cell Adhesion via Cas-Crk-C3G-Pathway-Mediated Rap1 Activation. Journal of Cell Science 2002, 115 (24), 4915–4924. 10.1242/jcs.00207.

(65) Cai, D.; Clayton, L. K.; Smolyar, A.; Lerner, A. AND-34, a Novel p130Cas-Binding Thymic Stromal Cell Protein Regulated by Adhesion and Inflammatory Cytokines. J Immunol 1999, 163 (4), 2104–2112.

(66) Sakakibara, A.; Hattori, S. Chat, a Cas/HEF1-Associated Adaptor Protein That Integrates Multiple Signaling Pathways. Journal of Biological Chemistry 2000, 275 (9), 6404–6410. 10.1074/jbc.275.9.6404.

(67) Bouton, A. H.; Riggins, R. B.; Bruce-Staskal, P. J. Functions of the Adapter Protein Cas: Signal Convergence and the Determination of Cellular Responses. Oncogene 2001, 20 (44), 6448–6458. 10.1038/sj.onc.1204785.

(68) Hitosugi, T.; Zhou, L.; Fan, J.; Elf, S.; Zhang, L.; Xie, J.; Wang, Y.; Gu, T.-L.; Alečković, M.; LeRoy, G.; Kang, Y.; Kang, H.-B.; Seo, J.-H.; Shan, C.; Jin, P.; Gong, W.; Lonial, S.; Arellano, M. L.; Khoury, H. J.; Chen, G. Z.; Shin, D. M.; Khuri, F. R.; Boggon, T. J.; Kang, S.; He, C.; Chen, J. Tyr26 Phosphorylation of PGAM1 Provides a Metabolic Advantage to Tumours by Stabilizing the Active Conformation. Nat Commun 2013, 4, 1790. 10.1038/ncomms2759.

(69) Makkinje, A.; Vanden Borre, P.; Near, R. I.; Patel, P. S.; Lerner, A. Breast Cancer Anti-Estrogen Resistance 3 (BCAR3) Protein Augments Binding of the c-Src SH3 Domain to Crk-Associated Substrate (P130cas). J Biol Chem 2012, 287 (33), 27703–27714. 10.1074/jbc.M112.389981.

(70) Moon, D. O. Deciphering the Role of BCAR3 in Cancer Progression: Gene Regulation, Signal Transduction, and Therapeutic Implications. Cancers (Basel) 2024, 16 (9), 1674. 10.3390/cancers16091674.

(71) Wiese, E. K.; Hitosugi, T. Tyrosine Kinase Signaling in Cancer Metabolism: PKM2 Paradox in the Warburg Effect. Front. Cell Dev. Biol. 2018, 6, 79. 10.3389/fcell.2018.00079.

(72) Chattopadhyay, A.; Vecchi, M.; Ji, Q.; Mernaugh, R.; Carpenter, G. The Role of Individual SH2 Domains in Mediating Association of Phospholipase C-Γ1 with the Activated EGF Receptor. Journal of Biological Chemistry 1999, 274 (37), 26091–26097. 10.1074/jbc.274.37.26091.

(73) Sakaguchi, K.; Okabayashi, Y.; Kido, Y.; Kimura, S.; Matsumura, Y.; Inushima, K.; Kasuga, M. Shc Phosphotyrosine-Binding Domain Dominantly Interacts with Epidermal Growth Factor Receptors and Mediates Ras Activation in Intact Cells. Molecular Endocrinology 1998, 12 (4), 536–543. 10.1210/mend.12.4.0094.

(74) Keilhack, H.; Tenev, T.; Nyakatura, E.; Godovac-Zimmermann, J.; Nielsen, L.; Seedorf, K.; Böhmer, F.-D. Phosphotyrosine 1173 Mediates Binding of the Protein-Tyrosine Phosphatase SHP-1 to the Epidermal Growth Factor Receptor and Attenuation of Receptor Signaling. Journal of Biological Chemistry 1998, 273 (38), 24839–24846. 10.1074/jbc.273.38.24839.

(75) Wang, P.; Zhou, R.; Zhou, R.; Feng, S.; Zhao, L.; Li, W.; Lin, J.; Rajapakse, A.; Lee, C.-H.; Furnari, F. B.; Burgess, A. W.; Gunter, J. H.; Liu, G.; Ostrikov, K. K.; Richard, D. J.; Simpson, F.; Dai, X.; Thompson, E. W. Epidermal Growth Factor Potentiates EGFR(Y992/1173)-Mediated Therapeutic Response of Triple Negative Breast Cancer Cells to Cold Atmospheric Plasma-Activated Medium. Redox Biol 2024, 69, 102976. 10.1016/j.redox.2023.102976.

(76) Saito, T.; Okada, S.; Ohshima, K.; Yamada, E.; Sato, M.; Uehara, Y.; Shimizu, H.; Pessin, J. E.; Mori, M. Differential Activation of Epidermal Growth Factor (EGF) Receptor Downstream Signaling Pathways by Betacellulin and EGF. Endocrinology 2004, 145 (9), 4232–4243. 10.1210/en.2004-0401.

(77) Batzer, A. G.; Rotin, D.; Ureña, J. M.; Skolnik, E. Y.; Schlessinger, J. Hierarchy of Binding Sites for Grb2 and Shc on the Epidermal Growth Factor Receptor. Mol Cell Biol 1994, 14 (8), 5192–5201. 10.1128/mcb.14.8.5192-5201.1994.

(78) Pelicci, G.; Lanfrancone, L.; Grignani, F.; McGlade, J.; Cavallo, F.; Forni, G.; Nicoletti, I.; Grignani, F.; Pawson, T.; Giuseppe Pelicci, P. A Novel Transforming Protein (SHC) with an SH2 Domain Is Implicated in Mitogenic Signal Transduction. Cell 1992, 70 (1), 93–104. 10.1016/0092-8674(92)90536-L.

(79) Okabayashi, Y.; Kido, Y.; Okutani, T.; Sugimoto, Y.; Sakaguchi, K.; Kasuga, M. Tyrosines 1148 and 1173 of Activated Human Epidermal Growth Factor Receptors Are Binding Sites of Shc in Intact Cells. Journal of Biological Chemistry 1994, 269 (28), 18674–18678. 10.1016/S0021-9258(17)32363-3.

(80) Palla, V.-V.; Karaolanis, G.; Katafigiotis, I.; Anastasiou, I.; Patapis, P.; Dimitroulis, D.; Perrea, D. Gamma-H2AX: Can It Be Established as a Classical Cancer Prognostic Factor? Tumour Biol. 2017, 39 (3), 101042831769593. 10.1177/1010428317695931.

(81) Perez, M.; García-Heredia, J. M.; Felipe-Abrio, B.; Muñoz-Galván, S.; Martín-Broto, J.; Carnero, A. Sarcoma Stratification by Combined pH2AX and MAP17 (PDZK1IP1) Levels for a Better Outcome on Doxorubicin plus Olaparib Treatment. Sig Transduct Target Ther 2020, 5 (1), 195. 10.1038/s41392-020-00246-z.

(82) Tu, W.-Z.; Li, B.; Huang, B.; Wang, Y.; Liu, X.-D.; Guan, H.; Zhang, S.-M.; Tang, Y.; Rang, W.-Q.; Zhou, P.-K. γH2AX Foci Formation in the Absence of DNA Damage: Mitotic H2AX Phosphorylation Is Mediated by the DNA-PKcs/CHK2 Pathway. FEBS Letters 2013, 587 (21), 3437–3443. 10.1016/j.febslet.2013.08.028.

(83) Kao, G. D.; Jiang, Z.; Fernandes, A. M.; Gupta, A. K.; Maity, A. Inhibition of Phosphatidylinositol-3-OH Kinase/Akt Signaling Impairs DNA Repair in Glioblastoma Cells Following Ionizing Radiation. Journal of Biological Chemistry 2007, 282 (29), 21206–21212. 10.1074/jbc.M703042200.

(84) Sourisseau, T.; Maniotis, D.; McCarthy, A.; Tang, C.; Lord, C. J.; Ashworth, A.; Linardopoulos, S. Aurora-A Expressing Tumour Cells Are Deficient for Homology-directed DNA Double Strand-break Repair and Sensitive to PARP Inhibition. EMBO Mol Med 2010, 2 (4), 130–142. 10.1002/emmm.201000068.

(85) Saki, M.; Makino, H.; Javvadi, P.; Tomimatsu, N.; Ding, L.-H.; Clark, J. E.; Gavin, E.; Takeda, K.; Andrews, J.; Saha, D.; Story, M. D.; Burma, S.; Nirodi, C. S. EGFR Mutations Compromise Hypoxia-Associated Radiation Resistance through Impaired Replication Fork–Associated DNA Damage Repair. Molecular Cancer Research 2017, 15 (11), 1503–1516. 10.1158/1541-7786.MCR-17-0136.

(86) Werner, A. N.; Kumar, A. I.; Charest, P. G. CRISPR-Mediated Reversion of Oncogenic KRAS Mutation Results in Increased Proliferation and Reveals Independent Roles of Ras and mTORC2 in the Migration of A549 Lung Cancer Cells. MBoC 2023, 34 (13), ar128. 10.1091/mbc.E23-05-0152.

(87) Korrodi-Gregório, L.; Soto-Cerrato, V.; Vitorino, R.; Fardilha, M.; Pérez-Tomás, R. From Proteomic Analysis to Potential Therapeutic Targets: Functional Profile of Two Lung Cancer Cell Lines, A549 and SW900, Widely Studied in Pre-Clinical Research. PLoS ONE 2016, 11 (11), e0165973. 10.1371/journal.pone.0165973.

(88) Donnelly, L. L.; Hogan, T. C.; Lenahan, S. M.; Nandagopal, G.; Eaton, J. G.; Lebeau, M. A.; McCann, C. L.; Sarausky, H. M.; Hampel, K. J.; Armstrong, J. D.; Cameron, M. P.; Sidiropoulos, N.; Deming, P.; Seward, D. J. Functional Assessment of Somatic *STK11* Variants Identified in Primary Human Non-Small Cell Lung Cancers. Carcinogenesis 2021, 42 (12), 1428–1438. 10.1093/carcin/bgab104.

(89) Tomas, A.; Futter, C. E.; Eden, E. R. EGF Receptor Trafficking: Consequences for Signaling and Cancer. Trends in Cell Biology 2014, 24 (1), 26–34. 10.1016/j.tcb.2013.11.002.

(90) Downward, J. Targeting RAS Signalling Pathways in Cancer Therapy. Nat Rev Cancer 2003, 3 (1), 11–22. 10.1038/nrc969.

(91) Stephen, A. G.; Esposito, D.; Bagni, R. K.; McCormick, F. Dragging Ras Back in the Ring. Cancer Cell 2014, 25 (3), 272–281. 10.1016/j.ccr.2014.02.017.

(92) Maziveyi, M.; Alahari, S. K. Cell Matrix Adhesions in Cancer: The Proteins That Form the Glue. Oncotarget 2017, 8 (29), 48471–48487. 10.18632/oncotarget.17265.

(93) Jiménez, C.; Portela, R. A.; Mellado, M.; Rodríguez-Frade, J. M.; Collard, J.; Serrano, A.; Martínez-A, C.; Avila, J.; Carrera, A. C. Role of the Pi3k Regulatory Subunit in the Control of Actin Organization and Cell Migration. The Journal of Cell Biology 2000, 151 (2), 249–262. 10.1083/jcb.151.2.249.

(94) Cain, R. J.; Ridley, A. J. Phosphoinositide 3-kinases in Cell Migration. Biology of the Cell 2009, 101 (1), 13–29. 10.1042/BC20080079.

(95) Dunne, P. D.; Dasgupta, S.; Blayney, J. K.; McArt, D. G.; Redmond, K. L.; Weir, J.-A.; Bradley, C. A.; Sasazuki, T.; Shirasawa, S.; Wang, T.; Srivastava, S.; Ong, C. W.; Arthur, K.; Salto-Tellez, M.; Wilson, R. H.; Johnston, P. G.; Van Schaeybroeck, S. EphA2 Expression Is a Key Driver of Migration and Invasion and a Poor Prognostic Marker in Colorectal Cancer. Clinical Cancer Research 2016, 22 (1), 230–242. 10.1158/1078-0432.CCR-15-0603.

(96) Miao, H.; Li, D.-Q.; Mukherjee, A.; Guo, H.; Petty, A.; Cutter, J.; Basilion, J. P.; Sedor, J.; Wu, J.; Danielpour, D.; Sloan, A. E.; Cohen, M. L.; Wang, B. EphA2 Mediates Ligand-Dependent Inhibition and Ligand-Independent Promotion of Cell Migration and Invasion via a Reciprocal Regulatory Loop with Akt. Cancer Cell 2009, 16 (1), 9–20. 10.1016/j.ccr.2009.04.009.

(97) Tang, Y.; Zang, H.; Wen, Q.; Fan, S. AXL in Cancer: A Modulator of Drug Resistance and Therapeutic Target. J Exp Clin Cancer Res 2023, 42 (1), 148. 10.1186/s13046-023-02726-w.

(98) De Gooijer, M. C.; Van Den Top, A.; Bockaj, I.; Beijnen, J. H.; Würdinger, T.; Van Tellingen, O. The G2 Checkpoint—a Node-based Molecular Switch. FEBS Open Bio 2017, 7 (4), 439–455. 10.1002/2211-5463.12206.

(99) Usatyuk, P. V.; Fu, P.; Mohan, V.; Epshtein, Y.; Jacobson, J. R.; Gomez-Cambronero, J.; Wary, K. K.; Bindokas, V.; Dudek, S. M.; Salgia, R.; Garcia, J. G. N.; Natarajan, V. Role of C-Met/Phosphatidylinositol 3-Kinase (PI3k)/Akt Signaling in Hepatocyte Growth Factor (HGF)-Mediated Lamellipodia Formation, Reactive Oxygen Species (ROS) Generation, and Motility of Lung Endothelial Cells. Journal of Biological Chemistry 2014, 289 (19), 13476–13491. 10.1074/jbc.M113.527556.

(100) Miyano, K.; Okamoto, S.; Yamauchi, A.; Kawai, C.; Kajikawa, M.; Kiyohara, T.; Tamura, M.; Taura, M.; Kuribayashi, F. The NADPH Oxidase NOX4 Promotes the Directed Migration of Endothelial Cells by Stabilizing Vascular Endothelial Growth Factor Receptor 2 Protein. Journal of Biological Chemistry 2020, 295 (33), 11877–11890. 10.1074/jbc.RA120.014723.

(101) Germon, Z. P.; Sillar, J. R.; Mannan, A.; Duchatel, R. J.; Staudt, D.; Murray, H. C.; Findlay, I. J.; Jackson, E. R.; McEwen, H. P.; Douglas, A. M.; McLachlan, T.; Schjenken, J. E.; Skerrett-Byrne, D. A.; Huang, H.; Melo-Braga, M. N.; Plank, M. W.; Alvaro, F.; Chamberlain, J.; De Iuliis, G.; Aitken, R. J.; Nixon, B.; Wei, A. H.; Enjeti, A. K.; Huang, Y.; Lock, R. B.; Larsen, M. R.; Lee, H.; Vaghjiani, V.; Cain, J. E.; De Bock, C. E.; Verrills, N. M.; Dun, M. D. Blockade of ROS Production Inhibits Oncogenic Signaling in Acute Myeloid Leukemia and Amplifies Response to Precision Therapies. Sci. Signal. 2023, 16 (778), eabp9586. 10.1126/scisignal.abp9586.

(102) Brown, D. I.; Griendling, K. K. Nox Proteins in Signal Transduction. Free Radical Biology and Medicine 2009, 47 (9), 1239–1253. 10.1016/j.freeradbiomed.2009.07.023.

(103) Kim, Y.-M.; Kim, S.-J.; Tatsunami, R.; Yamamura, H.; Fukai, T.; Ushio-Fukai, M. ROS-Induced ROS Release Orchestrated by Nox4, Nox2, and Mitochondria in VEGF Signaling and Angiogenesis. American Journal of Physiology-Cell Physiology 2017, 312 (6), C749–C764. 10.1152/ajpcell.00346.2016.

(104) Zulato, E.; Ciccarese, F.; Agnusdei, V.; Pinazza, M.; Nardo, G.; Iorio, E.; Curtarello, M.; Silic-Benussi, M.; Rossi, E.; Venturoli, C.; Panieri, E.; Santoro, M. M.; Di Paolo, V.; Quintieri, L.; Ciminale, V.; Indraccolo, S. LKB1 Loss Is Associated with Glutathione Deficiency under Oxidative Stress and Sensitivity of Cancer Cells to Cytotoxic Drugs and γ-Irradiation. Biochemical Pharmacology 2018, 156, 479–490. 10.1016/j.bcp.2018.09.019.

(105) Rosell, R.; Jantus-Lewintre, E.; Cao, P.; Cai, X.; Xing, B.; Ito, M.; Gomez-Vazquez, J. L.; Marco-Jordán, M.; Calabuig-Fariñas, S.; Cardona, A. F.; Codony-Servat, J.; Gonzalez, J.; València-Clua, K.; Aguilar, A.; Pedraz-Valdunciel, C.; Dantes, Z.; Jain, A.; Chandan, S.; Molina-Vila, M. A.; Arrieta, O.; Ferrero, M.; Camps, C.; González-Cao, M. KRAS-Mutant Non-Small Cell Lung Cancer (NSCLC) Therapy Based on Tepotinib and Omeprazole Combination. Cell Commun Signal 2024, 22 (1), 324. 10.1186/s12964-024-01667-x.

(106) Rizzo, S.; Sikorski, E.; Park, S.; Im, W.; Vasquez-Montes, V.; Ladokhin, A. S.; Thévenin, D. Promoting the Activity of a Receptor Tyrosine Phosphatase with a Novel pH-Responsive Transmembrane Agonist Inhibits Cancer-Associated Phenotypes. Protein Sci 2023, 32 (9), e4742. 10.1002/pro.4742.

(107) Perez-Riverol, Y.; Bandla, C.; Kundu, D. J.; Kamatchinathan, S.; Bai, J.; Hewapathirana, S.; John, N. S.; Prakash, A.; Walzer, M.; Wang, S.; Vizcaíno, J. A. The PRIDE Database at 20 Years: 2025 Update. Nucleic Acids Research 2025, 53 (D1), D543–D553. 10.1093/nar/gkae1011.

(108) Deutsch, E. W.; Bandeira, N.; Perez-Riverol, Y.; Sharma, V.; Carver, J. J.; Mendoza, L.; Kundu, D. J.; Wang, S.; Bandla, C.; Kamatchinathan, S.; Hewapathirana, S.; Pullman, B. S.; Wertz, J.; Sun, Z.; Kawano, S.; Okuda, S.; Watanabe, Y.; MacLean, B.; MacCoss, M. J.; Zhu, Y.; Ishihama, Y.; Vizcaíno, J. A. The ProteomeXchange Consortium at 10 Years: 2023 Update. Nucleic Acids Res 2023, 51 (D1), D1539–D1548. 10.1093/nar/gkac1040.

(109) Perez-Riverol, Y.; Xu, Q.-W.; Wang, R.; Uszkoreit, J.; Griss, J.; Sanchez, A.; Reisinger, F.; Csordas, A.; Ternent, T.; Del-Toro, N.; Dianes, J. A.; Eisenacher, M.; Hermjakob, H.; Vizcaíno, J. A. PRIDE Inspector Toolsuite: Moving Toward a Universal Visualization Tool for Proteomics Data Standard Formats and Quality Assessment of ProteomeXchange Datasets. Mol Cell Proteomics 2016, 15 (1), 305–317. 10.1074/mcp.O115.050229.

