## Supplementary Figures for "PTPRJ-Targeting Peptide Agonist Induces Broad Cellular Signaling Perturbations and DNA Damage in Lung Cancer Cells"

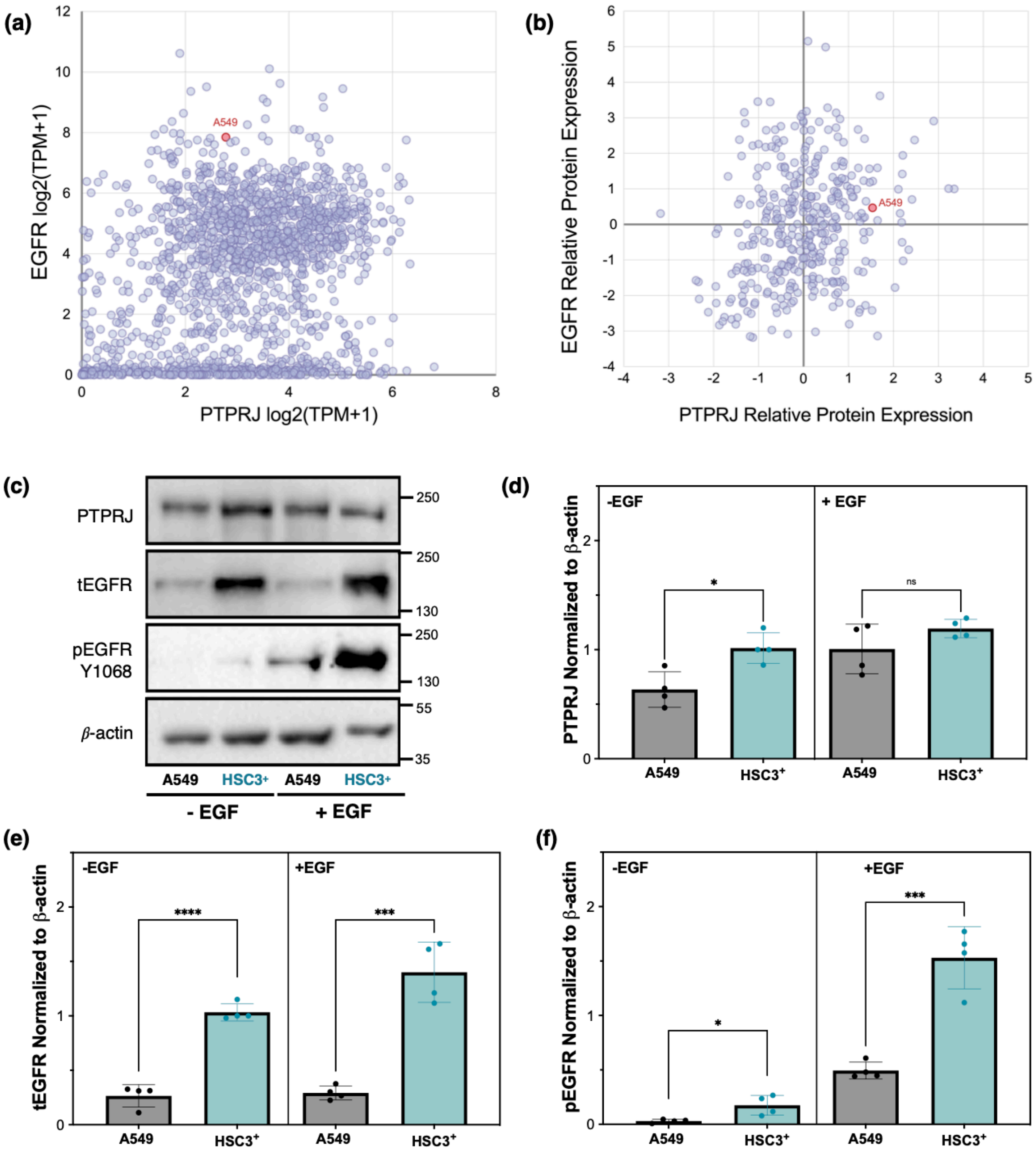


**Figure S1. (a)** Gene expression levels of *PTPRJ* and *EGFR*, and **(b)** relative protein expression of PTPRJ (Q12913) and EGFR (P00533) from harmonized MS (CCLE Gygi), DepMap Public 25Q3. **(c)** Representative immunoblot. Quantification of **(b)** total PTPRJ, **(c)** total EGFR, and **(d)** phosphorylated EGFR (Y1068) in A549 and HSC3 ectopically expressing PTPRJ (HSC3$^{+}$). The results are shown as the mean ± SD (n = 4). The statistical significance of the differences between samples was determined by one-way ANOVA with Tukey's multiple comparisons correction ($\alpha = 0.05$). ****, $p \leq 0.0001$; ***, $p \leq 0.001$; *, $p \leq 0.05$; ns, $p > 0.05$. Compared to the HSC3 cell line used in our earlier studies (engineered to express PTPRJ ectopically; HSC3$^{+}$), A549 shows a similar PTPRJ level but significantly lower EGFR levels. This difference in EGFR expression is also reflected in the higher basal and stimulated levels of phosphorylated EGFR (pY1068) in HSC3 cells.

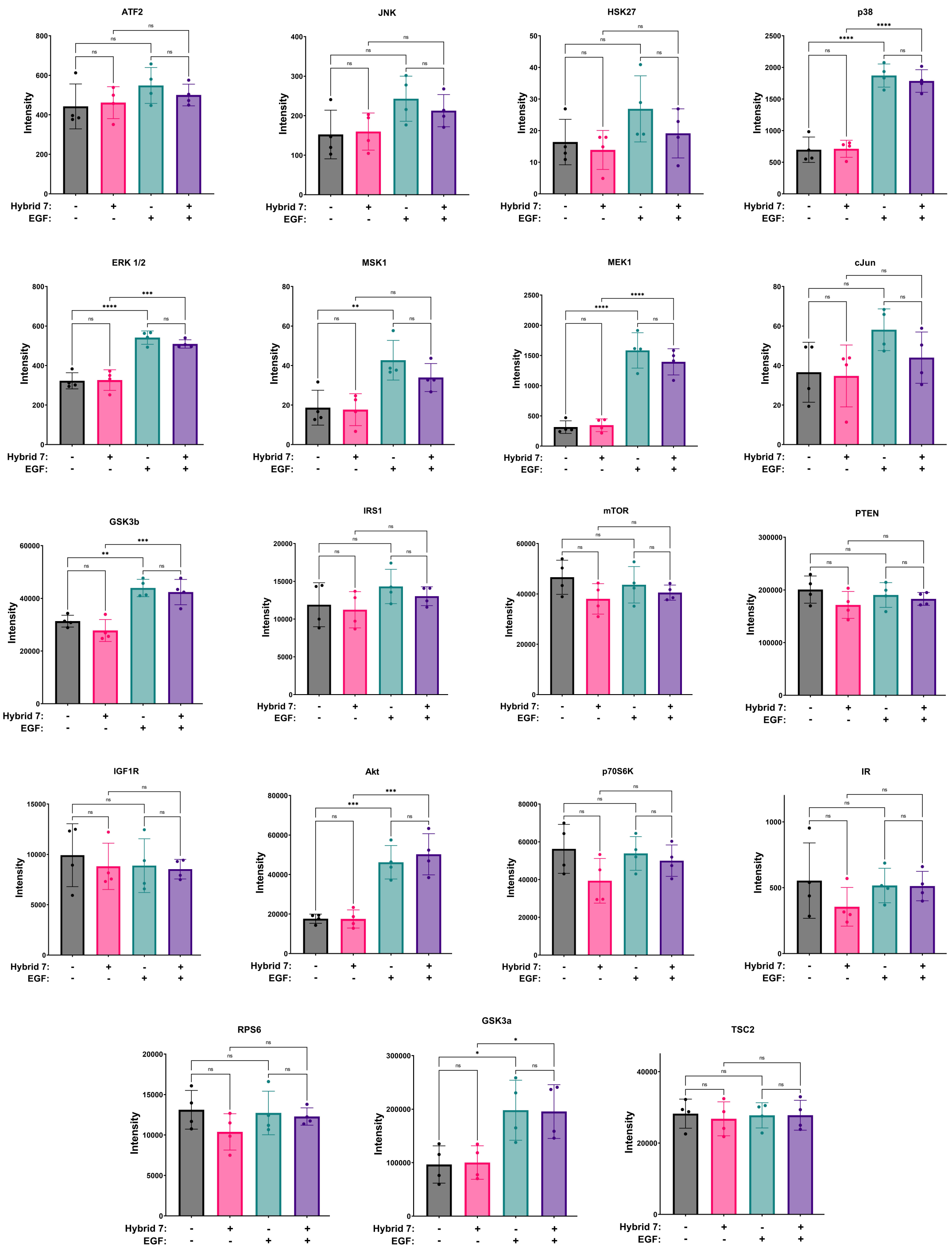


**Figure S2: Luminex of downstream signaling molecules in response to Hybrid 7**. A549 cells were treated with 10 µM Hybrid 7 for 5 minutes at 37°C, and pH was adjusted to pH 5.5 for 10 minutes at 37°C. After successive washes, cells were stimulated with EGF (10 ng/mL) for 10 minutes before lysis. Relative phosphoprotein abundances are shown as the mean ± SD (n=4). The statistical significance of differences between samples was determined by one-way ANOVA with Tukey's multiple comparisons correction (alpha=0.05). ****, $p \leq 0.0001$; ***, $p \leq 0.001$; **, $p \leq 0.01$; *, $p \leq 0.05$; ns, $p > 0.05$.

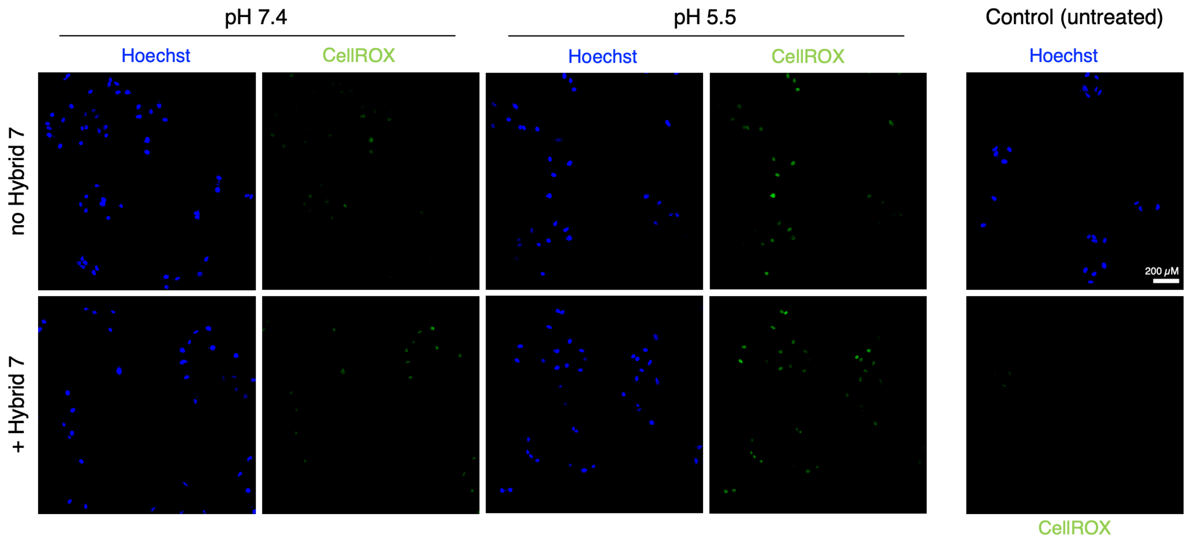


**Figure S3: Hybrid 7 increases ROS production.** Representative fluorescence images using the membrane-permeable CellROX Green reagent (5μM) are shown after a 10-minute treatment with Hybrid 7 (10 μM).
